# Rational Control of Basal CAR Expression Improves Discrimination in Inducible T Cell Circuits

**DOI:** 10.64898/2026.08.28.747722

**Authors:** Daniel Hoces, Jaclyn Ng, Julian Perez, Rogelio A. Hernández-López

## Abstract

SynNotch-CAR circuits improve T cell specificity by coupling antigen recognition to inducible CAR expression. However, basal CAR expression without receptor activation, termed here as leakiness, can reduce the separation between killing of intended target cells and sparing of antigen-positive off-target cells, limiting target-cell discrimination. Here, we systematically quantified basal CAR expression for several synNotch-CAR designs and developed a coupled ordinary differential equation model to show that discrimination depends on basal output, CAR potency, and effector-to-target ratio. We introduced C-terminal tags such as fluorescent proteins, degron domains, endocytosis signals, and endoplasmic reticulum retention motifs as a strategy to reduce CAR leakiness. We found that fluorescent proteins and degron-containing tags reduced basal CAR surface expression while preserving antigen-induced CAR expression, improving discrimination of antigen-density sensing and combinatorial circuits *in vitro*. In xenograft models, fluorescent protein-tagged CARs improved discrimination by reducing activity against off-target cells while retaining activity against high-antigen tumors. Degron-containing constructs reduced basal CAR expression *in vitro* but showed suboptimal performance *in vivo*, revealing a trade-off between basal CAR suppression and induced CAR persistence. Together, these findings demonstrate that basal output expression is a key parameter for inducible genetic circuit designs and establish layered transcriptional and post-translational regulation as a strategy to improve the fidelity of inducible T cell circuits.

## INTRODUCTION

Chimeric antigen receptor (CAR) T cell therapies have achieved remarkable clinical success in hematological malignancies, with multiple FDA-approved therapies now standard of care for patients with relapsed or refractory B cell cancers^1^. However, translating these successes to solid tumors has proven challenging, in part because many clinically relevant antigens, including HER2, EGFR, and GD2, are expressed on normal tissues, often at lower levels^2^. This antigen expression overlap creates a fundamental specificity challenge: conventional CAR T cells have limited ability to discriminate high-antigen expressing tumor cells from low-antigen expressing normal cells^3^. Here, we use *target-cell discrimination* to describe this functional separation between killing intended target cells (on-target) and sparing unintended target cells (off-target). This distinction is essential for safety because insufficient discrimination can result in on-target, off-tumor toxicity (OTOT) ^2,4–6^. OTOT has been observed in clinical trials targeting shared antigens, ranging from manageable cytokine release to life-threatening organ damage^3,7^. Addressing this specificity gap is therefore a central challenge for engineering safer and more precise T cell therapies for solid tumors.

To improve the specificity and programmability of engineered T cell recognition, several strategies have been developed, ranging from tuning individual CAR components, such as binding affinity and signaling strength^8,9^, to engineering receptor pairs or protein-based circuits that require simultaneous recognition of multiple antigens^10–13^. Among these approaches, inducible CAR circuits using synthetic Notch (SynNotch) receptors provide a distinct transcriptional-switch architecture in which antigen recognition is coupled to expression of a downstream output^14,15^. In SynNotch circuits, binding of a surface antigen triggers proteolytic cleavage of the receptor, releasing an orthogonal transcription factor that drives expression of a downstream effector, such as a CAR. This two-step architecture makes CAR expression and T cell cytotoxicity conditional on recognition of a priming antigen.

Two complementary strategies exploit this SynNotch-CAR circuit architecture to improve target cell discrimination (**Fig. 1A**). First, combinatorial dual-antigen circuits, in which a SynNotch receptor recognizes antigen A then induces a CAR targeting antigen B, thereby requiring both antigens for T cell killing ^16–18^. Second, antigen-density sensing circuits use tuned receptor affinities to distinguish high-antigen-density tumor cells from low-antigen-density normal cells. In this design, a low-affinity SynNotch receptor induces a high-affinity CAR for the same antigen, generating a sigmoidal dose-response that enables selective killing of high-density target cells while sparing low-density cells^19^. Together, these strategies establish SynNotch-CAR circuits as a versatile framework for engineering T cells with programmable target discrimination.

**Figure 1:**
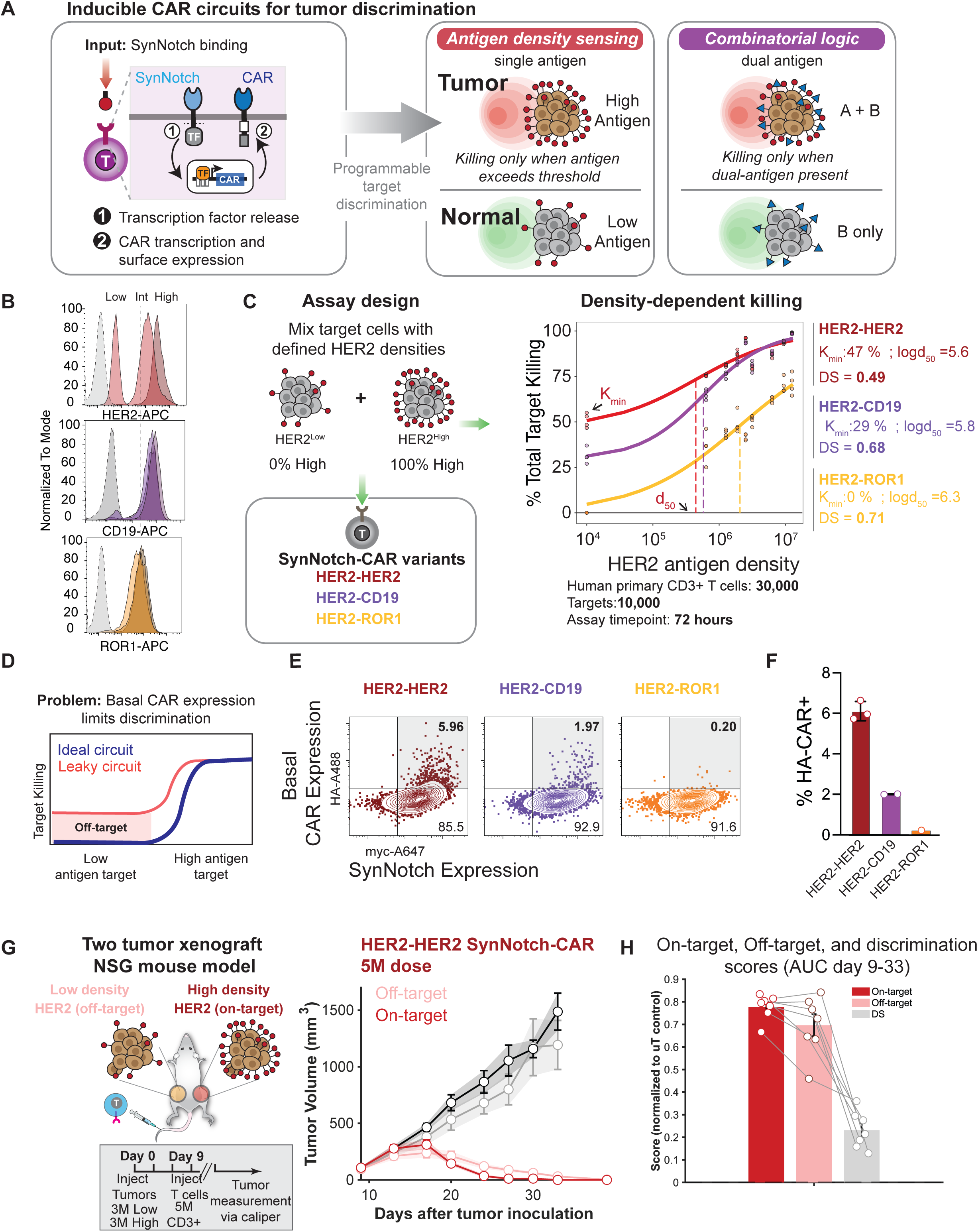
Basal CAR expression constrains discrimination in inducible SynNotch-CAR circuits. **(A) Left:** Schematic of SynNotch-to-CAR circuit for target discrimination. **Right:** Architectures to control CAR expression and killing via synNotch induction. Antigen density sensing uses a single target antigen and combinatorial logic a dual antigen recognition program. **(B)** Representative FACS histograms showing antigen density expression of engineered HCC1806 cell lines used in this study. The parental HCC1806 has low expression of HER2 (HER2^Low^) and ROR1, and no expression of CD19. Engineered HCC1806 cell lines express increasing levels of HER2 (HER2^Int^, HER2^High^). Parental and HER2 engineered cell lines were modified to express high levels of CD19 or ROR1. Isotype controls are shown in gray shades. **(C)** Schematic of *in vitro* killing assays to measure density dependent killing. HER2^Low^ target cells were mixed with HER2^Int^ or HER2^High^ target cells at different ratios to achieve a range of HER2 antigen densities. Mixed target cells were incubated with engineered T cells expressing an anti-HER2-SynNotch that induces different CAR variants (αHER2-BBz, αCD19-BBz, or αROR1-BBz) at a 3:1 effector to target ratio. Plot shows weighted average of HER2 molecules per cell in each mixed population vs. the percentage of targets killed after 72 hours. Each dot represents an individual replicate; curves were fitted to the data using a 4-parameter logistic curve. K_min_: percentage of low-antigen population killed by each circuit. logd_50_: log_10_ of weighted antigen density per cell at which 50% of the total population is killed by each circuit. DS: discrimination score. **(D)** Schematic depicting dose response of an ideal circuit (blue line) and a leaky circuit (red line) that displays off-target killing due to basal CAR expression. **(E-F)** Basal CAR expression in resting T cells (16 days post-activation). **(E)** Representative flow cytometry plots of synNotch and CAR expression as measured by epitope tags **(F)** %CAR+ (HA-Tag) T cells across independent experiments. **(G)** Schematic of two-tumor xenograft NSG mouse model to measure discrimination score *in vivo*. Mice were injected with HER2^Low^ (left flank) or mixed HER2^Low^:HER2^High^ (right flank, avg. HER2 density 10^6.9^ molecules/cell,) tumors. After 8 days, mice received 5 million of αHER2-SynNotch-to-αHER2-CAR or control CD3+ untransduced T cells. Plot shows tumor volume over time for HER2^Low^ (light lines) and HER2^High^ (dark lines) tumors with the HER2-HER2 circuit (red) or control T cell (black) groups. **(H)** On-target, Off-target and Discrimination score calculated using tumor volume areas under curve (AUC) from Day 9 to Day 33 (n=7/group). Individual data points represent single mice; bars represent mean ± SEM.

Despite these advances, most efforts to improve inducible T cell circuits have focused on optimizing the input sensing layer. Alternative SynNotch receptors have been developed to tune receptor activation, including designs that respond to distinct cell-cell tension ^20^. In parallel, alternative inducible promoters ^21,22^ and related modular receptors, such as synthetic intramembrane proteolysis receptors (SNIPRs), have expanded the range of inputs that can be coupled to customized gene-expression programs ^22,23^. These advances have improved the sensing fidelity and modularity of inducible T cell circuits. However, circuit performance depends not only on input sensing, but also on the output layer. In transcriptionally controlled circuits, low-level output gene expression can occur even in the absence of receptor activation, a phenomenon referred to as “leakiness”. When the output protein has a downstream signaling function, such as a CAR that activates T cells for killing, this basal expression may compromise the separation between uninduced and induced circuit states.

Basal or “leaky” CAR expression, the antigen-independent expression of CAR on the T cell surface, represents an important output-layer parameter in inducible CAR circuits. However, the functional consequences of basal CAR expression remain poorly defined. It is not known how much basal CAR expression can be tolerated before it reduces target-cell discrimination, whether this relationship differs across circuit designs, or how antigen density, effector-to-target ratio and T cell expansion together shape the resulting dose-response function. These questions are important because CAR receptors activate T cell for killing, even low levels of CAR might be sufficient to allow target recognition, T cell activation, proliferation, and cytotoxicity, thereby converting a small molecular signal into a measurable cellular response. Defining how basal CAR expression maps onto circuit discrimination is therefore essential for engineering inducible T cell circuits with predictable and robust function.

One general strategy to improve the separation between basal and induced circuit states is to layer transcriptional regulation with post-transcriptional or post-translational control ^24^. In this architecture, the input-sensing layer regulates when an output gene is transcribed, while an additional regulatory layer controls protein accumulation, trafficking, half-life, or activity at the cell membrane. Similar layered designs have been used in synthetic gene circuits and biosensors to suppress basal output, increase dynamic range, and sharpen input-output behavior ^25^. Whether this approach can successfully tune basal CAR expression while preserving antigen-induced activity has not been evaluated in inducible T cell circuits.

Here, we systematically characterize basal CAR expression across multiple SynNotch-CAR architectures, including antigen density sensing and combinatorial logic circuits, and quantify how leakiness affects target-cell discrimination across a range of antigen densities. We develop a mathematical model that couples CAR expression dynamics, T cell proliferation, and antigen density to predict circuit discrimination capacity. This framework identifies basal CAR expression as a critical but tunable parameter that constrains circuit fidelity across input designs. Guided by these quantitative insights, we evaluated post-translational control strategies in the form of C-terminal tags, including fluorescent proteins, degron domains, endocytosis signals and endoplasmic reticulum retention motifs, that reduce basal CAR surface expression. We validated these approaches in vitro and in vivo using single and dual-tumor xenografts, demonstrating improved discrimination in inducible CAR T cell circuits. Together, these results establish basal output expression as a quantitative design parameter for inducible T cell circuits and show that layered transcriptional and post-translational regulation can improve circuit fidelity by reducing basal activity while preserving antigen-induced function.

## RESULTS

### Basal CAR expression is associated with reduced discrimination in inducible SynNotch-CAR circuits

Inducible SynNotch-CAR circuits have been used to implement distinct recognition strategies, including antigen-density sensors and combinatorial dual-antigen logic gates that restrict CAR expression to cells expressing a defined priming antigen (**Fig.1A**). However, these circuit designs have not been systematically compared across antigen density inputs using a common quantitative framework. It remains unclear how recognition strategy, priming antigen density, and basal CAR expression determine the ability of inducible T cells to discriminate on-target from off-target cells. To address this question, we established an *in vitro* assay that evaluates the behavior of distinct synNotch-CAR circuit designs against target cell populations containing defined proportions of cells with low, intermediate, and high priming-antigen density. This approach enabled us to measure dose-response behavior and target cell discrimination across multiple circuit designs under matched experimental conditions.

We engineered target cells using the HCC1806 breast cancer cell line, which expresses HER2 at low levels (HER2^Low^, 10^4.2^ molecules/cell), within the average found in normal tissue (10^4.5^ molecules/cell)^19^. To evaluate antigen-density sensing, we generated cell lines expressing intermediate (HER2^Int^) and high (HER2^High^) HER2 densities. To evaluate combinatorial circuits, each target population was additionally engineered to express either CD19 (CD19^High^) or ROR1 (ROR1^High^) at high antigen density levels **(Fig. 1B)**. We then generated mixed target populations containing defined ratios of HER2^Low^ cells, HER2^Int^ cells, and HER2^High^ cells, producing a range of average HER2 densities that mimics heterogeneous tumor populations. Human primary CD3+ T cells were engineered with single lentiviral vectors encoding αHER2 (low affinity scFv) SynNotch→αHER2 (high affinity scFv) CAR (HER2-HER2)^19^, αHER2 SynNotch→αCD19 CAR (HER2-CD19), or αHER2 SynNotch→αROR1 CAR (HER2-ROR1) circuits, all CARs in this series had CD8TM and 41BB and CD3z costimulation and signaling domains. FACS sorted T cells were co-cultured with mixed target populations at an effector-to-target (E:T) ratio of 3:1 for 72 hours.

To establish a benchmark for target cell discrimination in this assay, we first evaluated conventional constitutively expressed αHER2 CAR T cells with either high- or low-affinity antigen recognition domains^19^ (**Fig. S2**). Consistent with our previous findings^19^, high-affinity and low-affinity CARs efficiently eliminated HER2^High^ target cells but also killed HER2^Low^ target cells. These results confirm that the HCC1806-based assay reproduces the discrimination problem previously observed for constitutive αHER2 CARs.

We next evaluated the dose response performance of the three inducible SynNotch-CAR circuits **(Fig. 1C)**. The HER2-HER2 and HER2-CD19 circuits efficiently responded to increasing HER2 density in the target population and eliminated most target cells at high antigen densities. The HER2-ROR1 circuit also responded to HER2 high-antigen density but was less efficient at eliminating the mixed target cell populations. To quantitatively compare inducible circuit performance, we calculated a discrimination score (DS), which integrates the ability of a circuit to eliminate cells with high antigen density (on-target cells) with sparing of cells expressing low-antigen levels (off-target), DS = 0 indicates no antigen-density discrimination; DS = 1 indicates perfect discrimination (see Methods for calculation). Despite their distinct designs, all three circuits exhibited suboptimal discrimination. HER2-HER2 circuits showed the lowest discrimination (DS = 0.49), followed by HER2-CD19 (DS = 0.65) and HER2-ROR1 (DS = 0.75) (**Fig. 1C** and **Fig. S3**). These discrimination scores deviate from perfect discrimination by distinct causes. HER2-HER2 and HER2-CD19 circuits eliminated a substantial fraction of off-target cells, killing 48% and 27.5% of HER2^Low^ target populations, respectively. In contrast, the HER2-ROR1 circuit showed no detectable killing of HER2^Low^ROR1^High^ off-target cells but was limited by reduced elimination of HER2^High^ROR1^High^ on-target targets. These results suggested that discrimination score can be compromised by at least two distinct factors: undesired killing of off-target cells or insufficient killing of on-target cells. We therefore sought to identify the circuit properties responsible for these differences in performance.

One potential explanation for the reduced discrimination due to off-target killing observed in HER2-HER2 and HER2-CD19 circuits is antigen-independent basal CAR expression, hereafter referred to as circuit leakiness **(Fig. 1D)**. To determine whether basal CAR expression was associated with discrimination performance, we quantified surface CAR expression in resting T cells (16 days after CD3/CD28 bead activation), prior to co-culture with target cells. We found differences in basal CAR expression across circuits. HER2-HER2 circuits showed the highest level of leakiness (∼6% CAR+ T cells), followed by HER2-CD19 (approximately 2%), whereas HER2-ROR1 circuits displayed minimal basal CAR expression (∼0.2%), as measured by an epitope HA-tag fused to the CAR construct **(Fig. 1E-F).** Similar %CAR+ results were observed when comparing HA-Tag staining with CD19-dextramer staining (**Fig. S6**). Remarkably, the rank order of basal CAR expression correlated with the discrimination profiles of the circuits, with HER2-HER2 circuits exhibiting both the highest leakiness and the lowest discrimination, whereas HER2-ROR1 circuits showed minimal leakiness and the highest discrimination score. Similar antigen-independent output expression was also observed in circuits built with αCD19-SynNotch and αCD19-SNIPR receptors, across a variety of CAR variants (**Fig. S1**), suggesting that basal CAR expression is not unique to αHER2 SynNotch circuits but may represent a more general property of transcriptionally regulated circuits.

To determine whether the CAR leakiness and suboptimal discrimination score observed *in vitro* would be important in a more complex setting, we established a two-tumor xenograft model in (NOD *scid* gamma) NSG mice. Mice were injected on contralateral flanks with HER2^Low^ and mixed HER2^High^/ HER2^Low^ cancer cells (the average HER2 density was 10^6.9^ molecules/cell) and treated with HER2-HER2 SynNotch-CAR T cells 8 days after tumor injection. Control T cells had no effect on either tumor, whereas HER2-HER2 circuit T cells reduced the volume of both tumors **(Fig. 1G)**. Thus, even modest levels of basal CAR expression (**Fig. S4A**) were sufficient to compromise discrimination *in vivo*, resulting in elimination of off-target cells expressing low HER2 density.

Taken together, these experiments demonstrate that transcriptionally regulated SynNotch-CAR circuits exhibit variable levels of leakiness. Across multiple circuit designs, basal CAR expression was associated with reduced discrimination and undesired elimination of off-target cells in both *in vitro* and *in vivo* settings. These findings identify basal CAR expression as an important factor affecting circuit fidelity and raise the question of how leakiness, target antigen density, and T cell killing efficacy collectively shape discrimination behavior.

### Mathematical model for SynNotch-CAR T cell function identifies basal CAR expression, killing potency and T cell dosage as key determinants of circuit discrimination

The T cell killing experimental results suggested that CAR leakiness can reduce the fidelity of inducible SynNotch-CAR circuits. However, the relationship between leakiness, antigen density and target cell discrimination is difficult to infer intuitively because CAR binding to its cognate antigen is coupled to T cell activation and proliferation. To quantitatively define how these parameters interact, we built a mathematical model of SynNotch-CAR T cell function. Using a system of coupled ordinary differential equations, we modeled target-cell killing, target-dependent T cell activation and proliferation, and CAR expression driven by either antigen-dependent SynNotch activation or antigen-independent basal output **(Fig. 2A, Fig. S7A-C)**. The model was parameterized using experimentally measured CAR expression to recapitulate the experimental dose-response killing curves (**Fig. 2B**). Key model outputs include off-target killing (K_min_), maximum on-target killing (K_max_), and the discrimination score (DS), which integrates both measurements into a quantitative metric of circuit performance.

**Figure 2.**
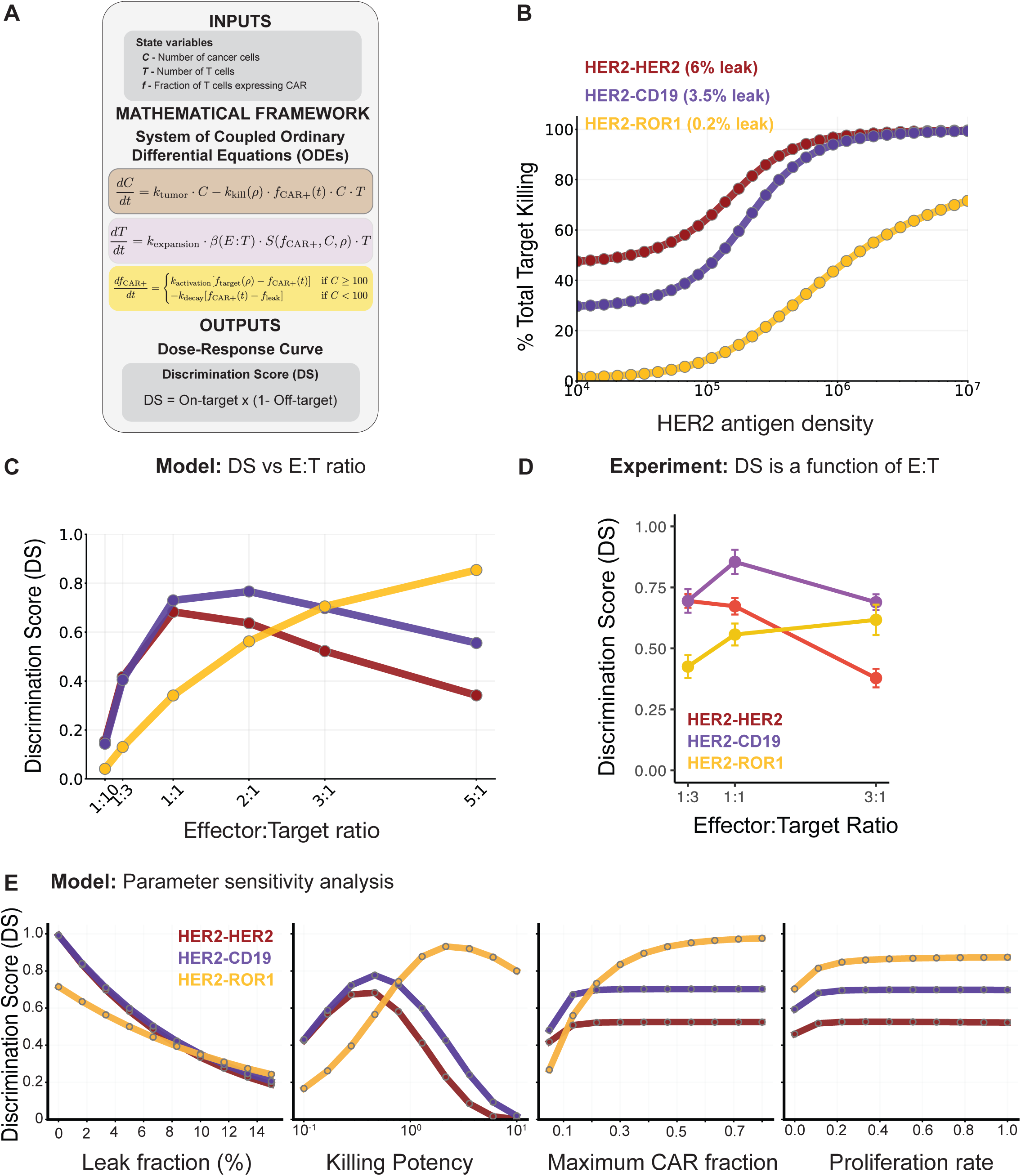
Mathematical modeling of synNotch-CAR T cell killing. **(A)** Summary of the system of non-linear ordinary differential equations (ODEs) model framework to describe SynNotch-CAR T cell killing activity as a function of antigen density. Initial variables include cancer cell count (*C*), T-cell count (*T*), and the fraction of T-cells expressing CAR (f_CAR+). The coupled system of ODEs integrates rates of tumor cell killing, antigen-dependent T-cell expansion, and synNotch-driven CAR induction to output dose-response curves and Discrimination Scores (DS = K_max_ * (1 − K_min_)) (See methods and Fig. S7 for more details). **(B)** Simulated percentage of total target killing after 72 hours of co-culture across HER2 antigen densities (10^4^ to 10^7^ molecules/cell) for three calibrated synNotch-CAR circuits with distinct baseline leak fractions: HER2-HER2 (6% leak), HER2-CD19 (3.5% leak), and HER2-ROR1 (0.2% leak). Curves show results for 30 simulated antigen densities. Model predictions **(C)** and experimental validation **(D)** of Discrimination Score (DS) as a function of Effector-to-Target (E:T) ratio. **(C)** Simulated DS across E:T seeding ratios (1:10 to 5:1) for HER2-HER2, HER2-CD19, and HER2-ROR1 SynNotch-CAR circuits. **(D)** Experimental DS evaluated across E:T ratios (1:3, 1:1, and 3:1) for the corresponding circuits. Data points represent mean ± SEM. (n=3) **(E)** Parameter sensitivity analysis on Discrimination Score (DS). Single-parameter sweeps evaluating the impact of model parameter variations on DS across all three constructs (HER2-HER2, HER2-CD19, and HER2-ROR1): Leakiness (leak fraction from 0% to 15%), Killing Potency (single-cell cytotoxicity multiplier from 0.1x to 10x on log scale), Max CAR Expression (maximal CAR expression fraction from 0.05 to 0.80), and Proliferation Rate (baseline expansion rate from 0.0 to 1.0).

We used the model to evaluate how different parameters affect circuit discrimination. The model predicted that discrimination depends on T cell dosage. At low effector to target (E:T) ratios, T cells are limiting, reducing overall target killing even at high antigen densities. At high E:T ratios, excess T cells amplify both intended on-target killing and unintended off-target killing, thereby reducing discrimination. Consequently, the model predicts an optimal E:T ratio at which the discrimination score (DS) is maximized, and that this optimum varies across circuit designs **(Fig. 2C, Fig. S7D)**. We tested this prediction experimentally by measuring the DS of HER2-HER2, HER2-CD19, and HER2-ROR1 synNotch-CAR circuits across different E:T ratios. Consistent with the model, each circuit displayed a distinct E:T dependent discrimination profile. The HER2-HER2 circuit achieved its highest DS at a lower E:T ratio (approximately 1:3), HER2-CD19 peaked near an E:T of 1:1, and HER2-ROR1 reached its highest DS at an E:T of 3:1 **(Fig. 2D)**.

We carried out a parameter sensitivity analysis and determined the DS as a function of CAR leakiness, killing potency, maximum fraction of CAR expressing T cells, and proliferation rate **(Fig. 2E, Fig. S8)**. CAR leakiness and CAR potency showed to have the largest impact, even a small fraction of CAR-expressing T cells initiates off-target killing and thus reduction of DS. As antigen density increases, SynNotch activation becomes the dominant source of CAR expression, leading to increased CAR+ T cell frequency, T cell expansion, and target cell killing (**Fig. S7E**). Thus, basal CAR expression compresses the dynamic range between the uninduced and induced states, thereby reducing discrimination. Similarly, the model predicted that there is a CAR potency that maximized the discrimination score. These results indicate that discrimination is not determined solely by circuit design, but emerges from the interaction between basal output expression, antigen induced CAR expression, CAR potency and T cell proliferation.

Together, the model and experimental validation establish basal CAR expression as a quantitative determinant of SynNotch-CAR circuit fidelity. Even low levels of leakiness can be amplified through T cell activation and proliferation, reducing the separation between killing of off- and on-target populations. These findings suggest that improving inducible circuit performance requires not only tuning the input-sensing layer but also controlling the abundance and potency of the CAR output.

### Two-layer regulation improves discrimination in inducible CAR circuits

The model suggested that reducing basal CAR expression should improve discrimination by increasing the separation between the uninduced and induced circuit activation states. To test this prediction, we designed a two-layer synNotch-CAR circuit by appending C-terminal regulatory tags to the CAR construct to impose an additional post-translational layer of control over CAR surface abundance (**Fig. 3A**). We engineered a series of two-layer, transcriptional and post-translational, HER2-HER2 SynNotch-CAR circuits and tested a panel of C-terminal motifs, including degron tags (cODC1 and variants^10,26^), endocytosis tails derived from CTLA-4^27^, endoplasmic reticulum retention signals^28–30^, and a fluorescent protein tag (mScarlet3). Among these, the fluorescent protein tag, and cODC1 degron reduced basal CAR expression without substantially compromising maximal induced CAR expression in a myc-bead activation assay **(Fig. 3B)**. Consistently, similar effects were observed across the two architectures, antigen density sensing and combinatorial dual-antigen circuits in experiments with high-antigen on-target cells (**Fig. 3C, D, Fig. S9A-B**).

**Figure 3.**
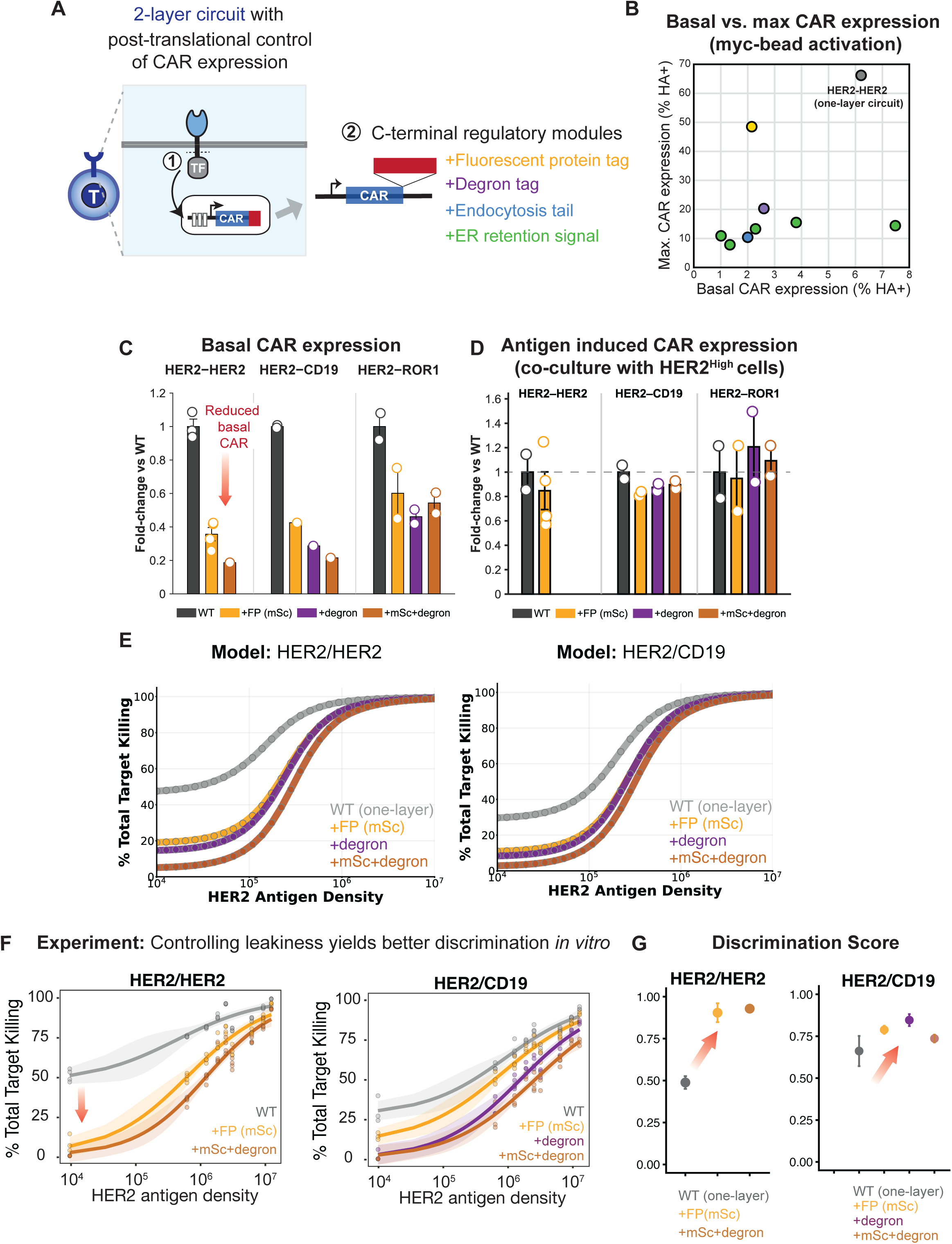
Post-translational control tunes basal CAR expression and target discrimination. **(A)** Schematics of a two-layer synNotch-CAR circuit. C-terminal regulatory tags were added to the CAR construct. **(B)** Scatter plot comparing basal CAR expression (% HA+) versus max CAR expression (max %HA+) across circuit variants, including one-layer HER2-HER2 circuit (WT, one-layer), and two-layer circuit variants that include C-terminal regulatory tags: fluorescent protein tag (+FP, mScarlet3, yellow), degron tag (+degron, mcODC, purple), endocytosis tags (CTL4-derived, blue) and endoplasmic reticulum retention tags (ERR, green). Basal surface CAR expression was measured 10 days post CD3/CD28 bead activation and max surface CAR expression was measured after 72 hours of co-culture with anti-myc magnetic beads. **(C)** Quantification of basal CAR expression fold-change relative to WT across HER2-HER2, HER2-CD19, and HER2-ROR1 synNotch-CAR circuits. **(D)** Quantification of antigen-induced CAR expression fold-change relative to WT in cocultures of circuit T cells with high-antigen density targets (10^7^ molecules/cell). **(E)** Model simulated dose response curves and **(F)** experimentally measured dose-response curves for HER2-HER2 (left panel) and HER2-CD19 (right panel) across one-layer circuit (WT, gray) and two-layer variants. **(G)** Discrimination Score (DS) quantification for HER2-HER2 (left) and HER2-CD19 (right) one-layer circuit (WT, gray) and different two-layer circuit variants. Circles indicate mean ± SEM (n =4 replicates).

To parameterize the effect of these post-translational motifs for the model, we quantified the fold-change in basal and antigen-induced CAR expression relative to the one-layer, transcriptional only, circuit. Across SynNotch-CAR designs, the mScarlet tag reduced basal CAR surface expression by approximately 60%, while degron-containing tags produced stronger reductions of approximately 3- to 5-fold depending on the circuit (**Fig. 3C, Fig. S9A**). In contrast, antigen-induced CAR expression was largely preserved for all the two-layer circuits across designs **(Fig. 3D, Fig. S9B)**. These measurements suggest that under the experimental conditions, these C-terminal tags preferentially reduce the uninduced output state while maintaining the induced state, thereby increasing the separation between basal and antigen-induced CAR expression. We used these experimentally measured fold-change values in basal CAR expression as inputs to the mathematical model to predict the behavior of the two-layer synNotch-CAR T circuits in target-cell killing and discrimination.

The model predicted that the observed reductions in basal CAR expression would markedly reduce off-target killing while largely preserving on-target killing. Specifically, reducing CAR leakiness was predicted to decrease off-target killing in HER2-HER2 circuits from approximately 45% to 5–15% in the two-layer circuits, and in HER2-CD19 circuits from approximately 30% to less than 10% (**Fig. 3E**). These predictions suggested that the second layer of regulation could improve circuit discrimination without eliminating the antigen-induced amplification required for effective killing of high-antigen on-target cells.

We experimentally validated these predictions in both antigen-density sensing and combinatorial logic circuits. For the HER2-HER2 circuits, addition of the fluorescent protein tag (+FP) or the combined fluorescent protein and degron (+mSc+degron) reduced killing of off-target cells while preserving killing of on-target cells **(Fig. 3F, left)**. Similar improvements were observed for HER2-CD19 circuits with the fluorescent protein (+FP), degron (+degron), and their combination (+mSc+degron) **(Fig. 3F, right)**. Although some variants showed a modest reduction in maximal killing of high-antigen on-target cells, most two-layer circuits achieved a marked improvement in discrimination score relative to the one-layer counterparts. For the HER2-HER2 circuits, the DS increased from 0.49 in the one-layer circuit to 0.90 with +FP and 0.93 with +mSc+degron **(Fig. 3G)**. However, when the one-layer circuit showed low basal leakiness, as in the case of another HER2-ROR1 circuit (ScFv: Clone F, CD28-CD3z) (**Fig. S9D**), the DS score for the one-layer circuit was high, and the C-terminal CAR modifications did not have a major impact in their function **(Fig. S9E-F).**

The observed improvement in discrimination score for the two-layer circuits was primarily driven by reduced surface CAR expression at baseline. The two-layer synNotch-CAR circuits showed similar percentages of CAR+ cells and similar T cells numbers compared with the one-layer HER2-CD19 circuit in co-cultures with targets expressing different HER2 densities **(Fig. S9C)**. Thus, post-translational control of the CAR output reduced basal CAR activity without eliminating the antigen-induced amplification required for effective target-cell killing.

We next tested whether addition of post-translational output regulation improved discrimination *in vivo* using the two-tumor xenograft model described above. Low- and high-antigen density tumors were established on contralateral flanks of NSG mice, and mice received 5 million engineered T cells. In contrast to the one-layer HER2-HER2 circuit, which reduced the growth of both HER2^Low^ and HER2^High^ tumors (**Fig. 1G**), the two-layer HER2-HER2+mScarlet showed low levels of CAR leakiness (**Fig. S11A**) and selectively reduced the volume of the HER2^High^ tumor while allowing continued growth of the HER2^Low^ tumor **(Fig. 4A, Fig. S11B)**. In contrast to the *in vitro* results, adding the C-terminal degron tag or a combined mScarlet+degron tag showed lower discrimination score than the mScarlet tag *in vivo* (**Fig. 4B**).

**Figure 4:**
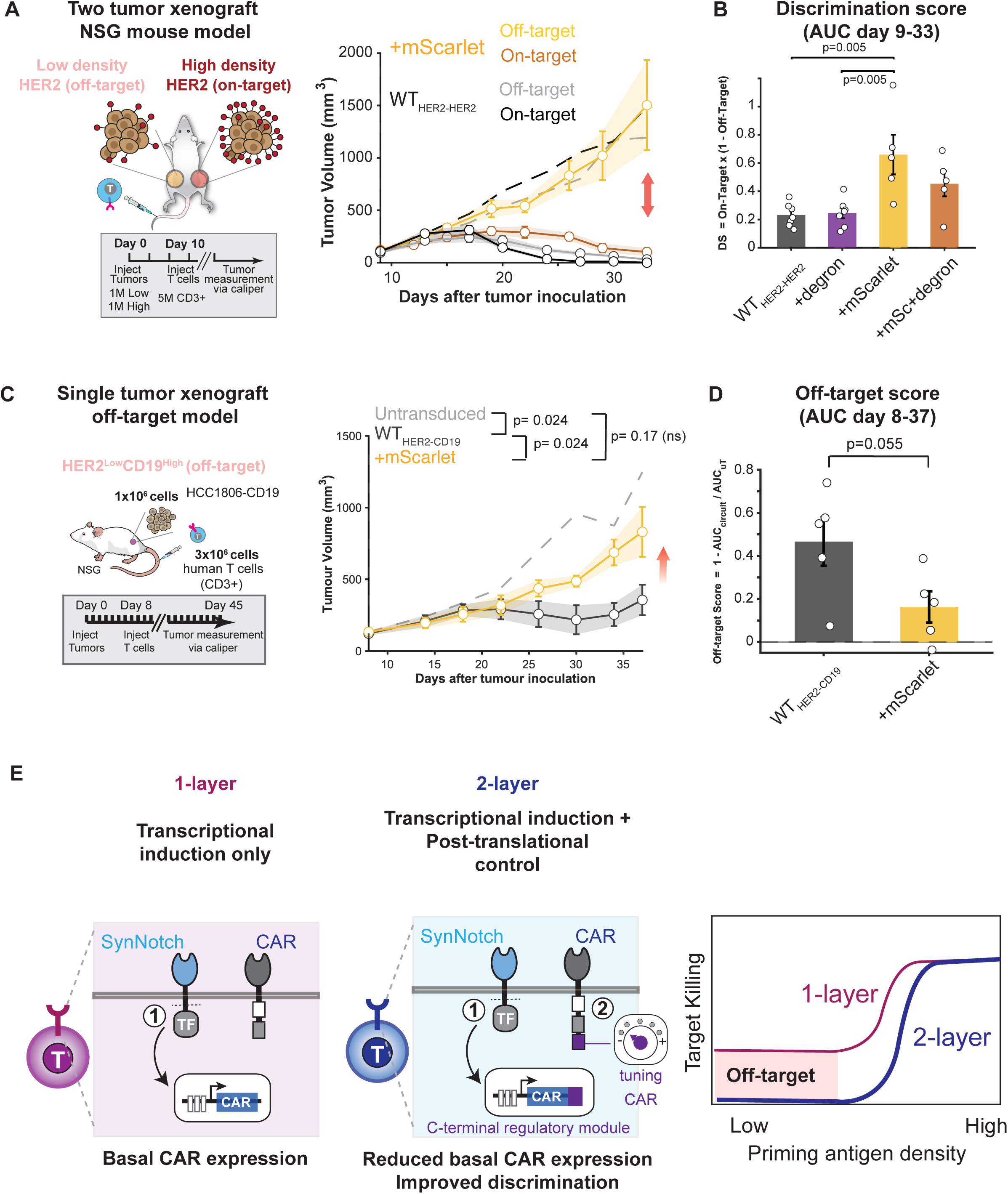
Two-layer synNotch-CAR circuits with post-translational control improve *in vivo* discrimination. **(A)** Schematic of two-tumor xenograft NSG mouse model to measure discrimination score: 1 million HER2^low^ HCC1806 (off-target, left flank) and 1 million mixed HCC1806 (25% HER2^low^/75% HER2^high^) (on-target, right flank) were implanted in NSG mice. Tumor volumes of mice treated with 5 million control CD3+ T cells (n=7), WT HER2-HER2 T cells (n=7), or HER2-HER2+mScarlet T cells (n=5). **(B)** Discrimination score calculated using tumor volume areas under curve from Day 9 to Day 33 of various treatment groups: WT HER2-HER2 T cells (n=7), HER2-HER2+degron T cells (n=7), HER2-HER2+mScarlet T cells (n=5), or HER2-HER2+mSc+degron T cells (n=5). Individual data points represent single mice; bars represent mean ± SEM. Group comparisons were performed using the Mann-Whitney U test with Bonferroni correction for multiple comparisons. **(C)** Schematic of single-tumor xenograft NSG mouse model to measure off-target score: 1 million of HER2^low^ CD19^high^ HCC1806 cells were implanted into the right flank of NSG mice and treated with T cells 8 days after tumor inoculation. Tumor volumes of mice treated with 3 million control CD3+ T cells, WT HER2-CD19 T cells, or HER2-CD19+mScarlet T cells (n=5/group). **(D)** Off-target score calculated based on tumor volume area under curve (AUC) from Day 8 to Day 37 of WT HER2-CD19 and HER2-CD19+mSc (n=5/group) treatment groups. Individual data points represent single mice; bars represent mean ± SEM. Group comparisons were performed using the Mann-Whitney U test. See Figures S11-S17 for individual tumor data and additional tumor models. **(E)** Schematic of single layer transcriptional control circuit and two-layer, transcriptional and post-translational control, circuits and their corresponding dose-response target killing curves.

To determine the function of the two-layer combinatorial logic circuits, we evaluated the HER2-CD19 SynNotch-CAR circuits in a single-tumor xenograft model using HER2^Low^CD19^High^ target cells. In this setting, target cells express the CAR antigen CD19 at high density but only low levels of the priming HER2 antigen, providing a test for any basal CAR-dependent activity in the absence of a strong SynNotch activation. The one-layer HER2-HER2 and one-layer HER2-CD19 circuits controlled HER2^Low^CD19^High^ tumor growth, consistent with their leaky αHER2 CAR and αCD19 CAR expression (**Fig. S12A**) driving off-target activity (**Fig. 4C**, and **Fig. S12B**). In contrast, addition of the mScarlet tag or a degron tag to the HER2-CD19 circuit reduced CAR leakiness (**Fig. S12A**) and tumor control with growth curves similar to those observed in mice treated with untransduced T cell controls (**Fig. 4C,D and Fig. S12-S13**).

Additional *in vivo* experiments with HER2-CD19 and another HER2-ROR1 SynNotch-CAR circuit variants further supported the idea that post-translational output control can modulate circuit behavior to improve discrimination, although the magnitude of the effect depended on circuit variant and experimental context. In a dual-tumor HER2^Low^CD19^High^/HER2^High^CD19^High^ model, the two-layer mScarlet-tagged HER2-CD19 circuit showed a trend toward improved discrimination, although non-significant when compared to the one-layer circuit, but tumor control was incomplete (**Fig. S14**).

We also tested the one- and two-layer circuits in a HER2^High^ROR1^High^ xenograft model, taking advantage of a previously established model of on-target off-tumor toxicity. In this model we used the Clone F of the αROR1 antibody, which cross-reacts with murine ROR1^31^, to build synNotch-CAR circuits. The one-layer synNotch-CAR circuit showed ∼10% leaky CAR expression (**Fig. S15A**), however this did not result in detectable off-target killing *in vitro*. In co-cultures with HER2^Low^ROR1^High^ (off-target) and HER2^High^ROR1^High^ (on-target) engineered lines, the synNotch-CAR achieved significantly higher discrimination scores than the constitutive CAR across multiple E:T ratios (**Fig. S15B-C**).

When tested in tumor-free mice at a high dose (8 million T cells), the one-layer SynNotch-CAR only delayed the fatal toxicity observed for the constitutive CAR, suggesting that leaky expression was sufficient to drive off-target killing at this dose (**Fig. S16**). This toxicity was dose dependent: reducing the dose to 4 million T cells eliminated apparent toxicity for both the constitutive and SynNotch-CAR circuit. We therefore reasoned that a comparable T cell dose could enable synNotch-CAR circuits to target HER2^High^ROR1^High^ tumors while sparing tissues that express lower ROR1 levels.

To test the on-target function of HER2-ROR1 synNotch-CAR T cell circuits, we engineered a one-and two-layer circuits with a C-terminal fluorescent protein tag. Both circuits showed minimal leaky CAR expression (**Fig. S17A**). At 5 million T cell dose, one- and two-layer mScarlet-tagged circuits achieved comparable on-target tumor control relative to untransduced T cells (**Fig. S17B-D**), with no significant difference in on-target score between the one- and two-layer circuits. These data indicate that basal expression is not sufficient by itself to predict circuit behavior; its functional impact depends on the potency and persistence of the induced CAR output.

### CAR surface persistence may contribute to the *in vivo* tradeoff between basal-output suppression and induced function

In vitro, degron-containing circuits improved discrimination relative to the one-layer circuits by reducing basal CAR surface expression and limiting off-target killing. However, this reduction in basal output did not correlate consistently with improved performance *in vivo*. In both the HER2-CD19 single-tumor off-target-killing model (**Fig. S12**) and the HER2-HER2 dual-tumor discrimination model (**Fig. S11**), degron-containing circuits showed less discrimination than the mScarlet-tagged circuit. These results suggested that reducing basal CAR expression is not sufficient to optimize circuit performance, because the same mechanisms that suppress basal CAR expression may also reduce the duration of induced CAR availability after SynNotch priming, potentially compromising long-term killing. To test this possibility, we compared post-induction killing dynamics following SynNotch activation with α-myc beads (**Fig. S10B**). The mScarlet-tagged circuit sustained killing activity similarly to the unmodified circuit after varying resting periods, whereas degron- and endocytosis-tagged circuits lost killing activity more rapidly (**Fig. S10C,D**). These results indicate that optimal output-layer regulation requires balancing leakiness suppression with enough abundance and persistence of antigen-induced CAR.

Together, these results demonstrate that basal output expression is a critical determinant of inducible circuit behavior and that layered transcriptional and post-translational regulation provides a general strategy to improve discrimination fidelity. The two-layer design, transcriptional induction and post-translational control, preserves the ability of SynNotch-CAR circuits to amplify responses to high-antigen on-target cells while reducing inappropriate activity in low-antigen off-target contexts (**Fig. 4E**). These findings establish output-layer regulation as an independent design axis for engineering inducible CAR circuits with improved fidelity.

## DISCUSSION

A central challenge of inducible circuits is that even small amounts of basal output can produce substantial unintended biological effects. Here we show that basal CAR expression limits the fidelity of SynNotch-CAR circuits designed to discriminate target cells and that layering post-translational regulation on top of transcriptional control substantially improves discrimination across both antigen-density sensing and combinatorial logic architectures.

We show that even low levels of basal CAR expression (**Fig. 1E**), are enough for off-target recognition and killing. In this study, we used a single vector to encode both the synNotch receptors and the inducible promoter that controls the CAR and showed that leakiness was present across a variety of circuit variants (**Fig. 1E, Fig. S1**). Interestingly, there seems to be a correlation between the CAR identity and the degree of leakiness. Even when all circuits express the same HER2 SynNotch, they still differ in the observed leakiness depending on the induced CAR. Further studies would be needed to elucidate the molecular basis of this behavior. Nevertheless, small differences in circuit leakiness and therefore CAR expression can result in substantial functional killing differences, both *in vitro* and *in vivo*. By quantifying this through a discrimination score and dose-response curves across multiple antigen densities, we provide a framework for comparing circuit designs and identifying performance limits.

To understand how basal CAR expression, antigen-dependent induction, CAR potency, and T cell proliferation determine population-level circuit performance, we developed a mathematical framework based on coupled ordinary differential equations (**Fig. 2A**). Predicting the dose-response behavior of these inducible circuits is difficult because small basal CAR+ populations can expand upon target engagement, converting negligible baseline signals into substantial killing. Our model captured these factors and predicted dose-response curves and discrimination scores that match experimental measurements.

Mathematical models have been increasingly used to describe CAR T cell killing dynamics, including predator-prey frameworks that capture dose-dependent tumor killing, T cell proliferation, and exhaustion^32^, as well as data-driven approaches that infer functional response classes from time-series cytotoxicity data^33^. More closely related to our study, PASCAR provides a multiscale framework for constitutive and SynNotch-inducible CAR T cells by integrating receptor-ligand binding, single-cell variability in CAR and ligand abundance, kinetic proofreading, population dynamics, and Pareto optimization of CAR affinity and SynNotch threshold parameters^34^. Our model is complementary to these approaches. Rather than building a high-resolution description of receptor signaling or single-cell variability, we focused on a minimal set of coupled dynamics needed to isolate basal CAR expression as an explicit design parameter. This simplified formulation allowed us to test how leakiness interacts with antigen density, CAR potency, T cell expansion, and effector-to-target ratio to shape target-cell discrimination. In this framework, even small basal CAR-positive fractions can generate substantial off-target killing when coupled to T cell activation and expansion. Future work could integrate this leakiness parameterization into higher-resolution models such as PASCAR, combining single-cell variability with basal-output control to more completely predict the fidelity of inducible CAR circuits.

Layering regulatory mechanisms is a common strategy in synthetic biology to improve the fidelity of synthetic circuits^24,35–37^. Transcriptional regulation determines when a gene is expressed but often incompletely suppresses basal protein accumulation. Rather than relying exclusively on increasingly tight transcriptional control, many synthetic systems combine transcriptional regulation with post-transcriptional or post-translational mechanisms to suppress basal activity while preserving robust induction^24,25^. Similar strategies have improved performance in bacterial metal biosensors, metabolic circuits, and mammalian gene circuits^38–40^.

We apply this principle to genetically encoded inducible CAR circuits in human primary T cells. Post-translational control substantially improved circuit discrimination while preserving robust induction. We tested this systematically by appending C-terminal regulatory modules to the CAR: fluorescent proteins, degron tags, endocytic tails, ER-retention signals and combinations. Some of these modular elements successfully reduced basal CAR surface expression and improved discrimination in both antigen-density-sensing and combinatorial dual-antigen circuits, while maintaining robust antigen-induced CAR expression. This suggests a general design principle: transcriptional control determines when the circuit activates; post-translational control determines how much output accumulates.

A key finding in this study is that reducing basal CAR expression alone is not sufficient to optimize circuit performance. Although degron-containing constructs, specifically the +degron and +mScarlet+degron variant, showed improved discrimination in vitro (similar to +mScarlet) (**Fig. 3F**) they show suboptimal behavior *in vivo* relative to the mScarlet-tagged circuit in the dual-tumor HER2-CD19 and HER2-HER2 on-target/off-target models (**Fig. 4 and Fig. S14**). These results suggest that the same mechanism that reduces basal CAR expression may also shorten the duration of the CAR at the surface after induction, thereby limiting sustained tumor control. In contrast, the mScarlet fusion reduced basal CAR expression while retaining sufficient antigen-induced expression and functional persistence to improve discrimination in vivo. The ER retention tails had a similar CAR surface decay behavior that the +mScarlet and WT circuit but did not produce enough surface CAR upon induction. Thus, the optimal output-control module is not necessarily the one that produces the lowest basal CAR expression, but rather the one that achieves the most favorable balance between suppressing leakiness and preserving sufficient CAR abundance and persistence after antigen-dependent induction. Additional experiments measuring CAR surface abundance and functional activity over time will be important for defining the mechanisms underlying these differences.

The translational implementation of this strategy will require regulatory modules that are compatible with clinical cell therapies. Although the fluorescent protein mScarlet provided a useful proof-of-principle output-control module, fluorescent proteins are non-human sequences and may create immunogenicity concerns in clinical applications. The relevant design principle, however, is independent of the specific fluorescent protein used here. Future work can use the quantitative framework developed in this study to identify alternative regulatory elements, including minimal degrons, trafficking motifs, endocytosis modules, and inducible stabilization strategies, that achieve a similar balance between basal-output suppression and induced CAR persistence. These efforts may also be combined with optimization of promoter strength, transcription-factor activity, receptor affinity, or SynNotch design to create multilayer circuits tailored to specific target antigens and therapeutic contexts.

Our results also show that the functional consequences of basal CAR expression depend on the potency of the downstream CAR and the effective T cell dose. The high-potency αHER2 CAR (high affinity scFv, 41BB-CD3z) showed strong off-target killing at a 3-5M dose in a single- and dual-tumor models, making leakiness effects readily detected *in vivo*. In contrast, the αROR1 CAR (F clone scFv-CD28-CD3z ITAM mutant) required much higher T cell doses (8 million cells), before the effect of circuit leakiness became apparent **(Fig. S16A**). These findings illustrate that similar degrees of basal CAR expression can have distinct functional consequences depending on receptor signaling strength, cytotoxic potency, T cell expansion, and target-cell antigen density abundance. They also underscore the value of evaluating inducible circuits across antigen-density gradients and dose ranges, because basal output that appears inconsequential in one assay condition may become functionally important under conditions of increased T cell dosage, prolonged antigen exposure, or more potent CAR signaling.

Several important questions remain. Our work focused on SynNotch-CAR circuits targeting defined antigen densities in xenograft models. Future studies should test performance in more complex tumor microenvironments, for example antigen expression changing over time. It will also be important to define how the circuit behavior aligns with antigen-expression distributions in patients’ tumors and candidate normal tissues, which may ultimately help guide target selection, circuit design, and patient stratification.

In summary, we identify basal CAR expression as a quantitative determinant of inducible circuit fidelity and demonstrate that layering post-translational regulation onto transcriptional control can improve discrimination. More broadly, our work establishes a general design principle for engineering inducible therapeutic circuits: when low levels of basal output can be functionally amplified, controlling transcription alone may be insufficient. Combining quantitative measurements, mathematical modeling, and modular circuit engineering provides a framework for designing higher-fidelity T cell circuits across a wide range of synthetic biology applications.

## Acknowledgments

We thank members of the Hernandez-Lopez Lab for advice and helpful discussions; C. Prange, J. Miguens and Q. Xue for critical reading of this manuscript. C. Mackall for sharing DNA plasmids; the Stanford shared FACS and Veterinary service core facilities for assistance, and advice with data collection.

## Author contributions

Conceptualization: D.H., J.N. J.P, R.A.H.-L

Formal analysis: D.H., J.N. J.P, R.A.H.-L

Funding acquisition: R.A.H.-L.

Investigation: D.H., J.N. J.P

Methodology: D.H., J.N. J.P

Project administration: R.A.H.-L.

Supervision: D.H., R.A.H.-L.

Visualization: D.H., J.N. J.P, R.A.H.-L

Writing – original draft: D.H., J.N. J.P, R.A.H.-L

Writing – review & editing: D.H., J.N. J.P, R.A.H.-L

---

**Figure S1:**
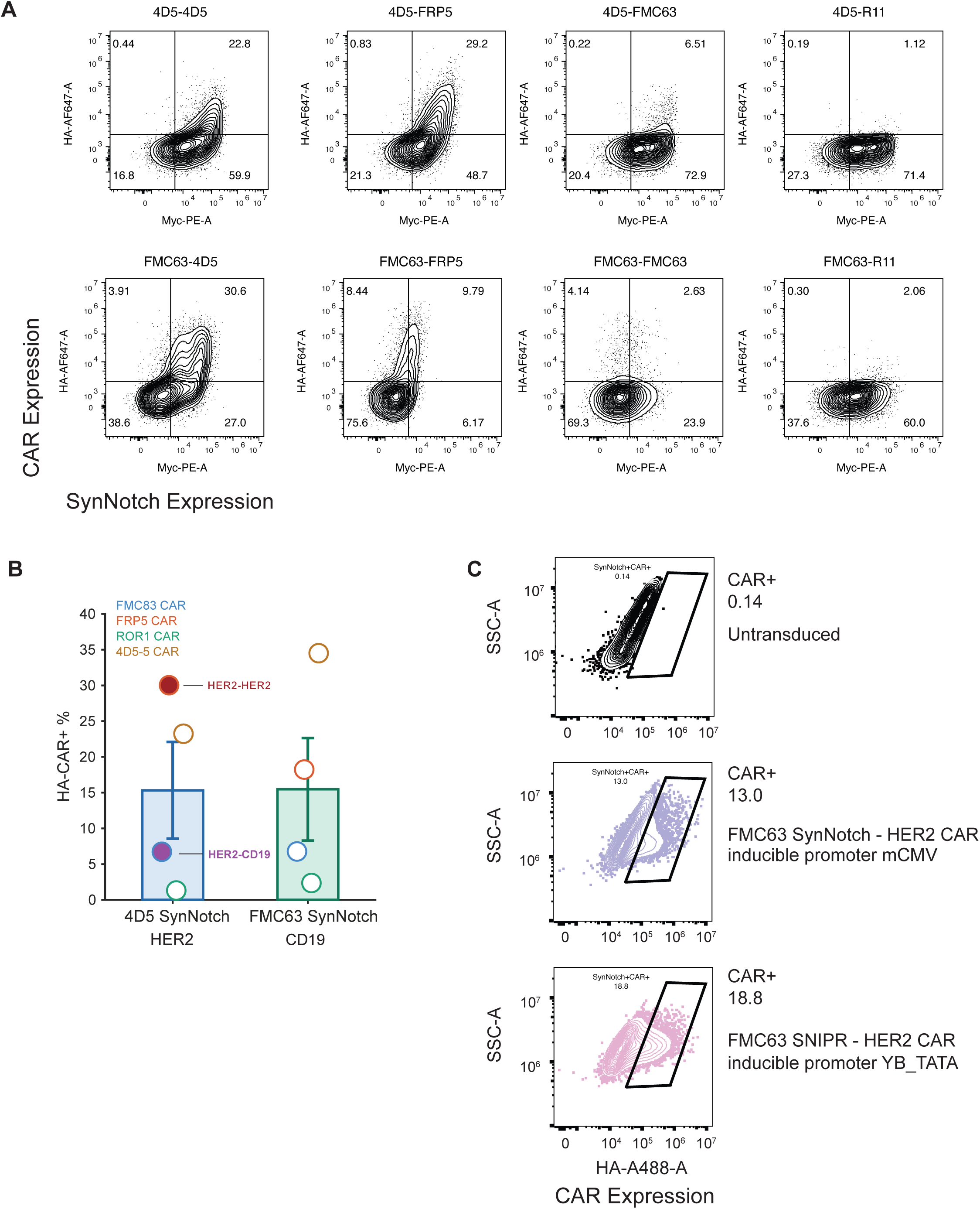
Basal CAR expression across representative SynNotch-CAR and SNIPR-CAR circuits. **(A-B)** SynNotch and basal CAR expression in resting T cells across different combinations of SynNotch receptors (anti-HER2 4D5 Low affinity scFv, anti-CD19 FMC63 scFv) and CAR constructs (anti-HER2 4D5 scFv and FRP5 scFv, anti-CD19 FMC63 scFv and anti-ROR1 R11 scFv). **(A)** Representative flow plots and **(B)** basal CAR expression summary plot. **(C)** Representative flow plots comparing basal CAR expression (%HA+) in anti-CD19 SynNotch vs anti-CD19 SNIPR circuits both with an anti-HER2 CAR.

**Figure S2:**
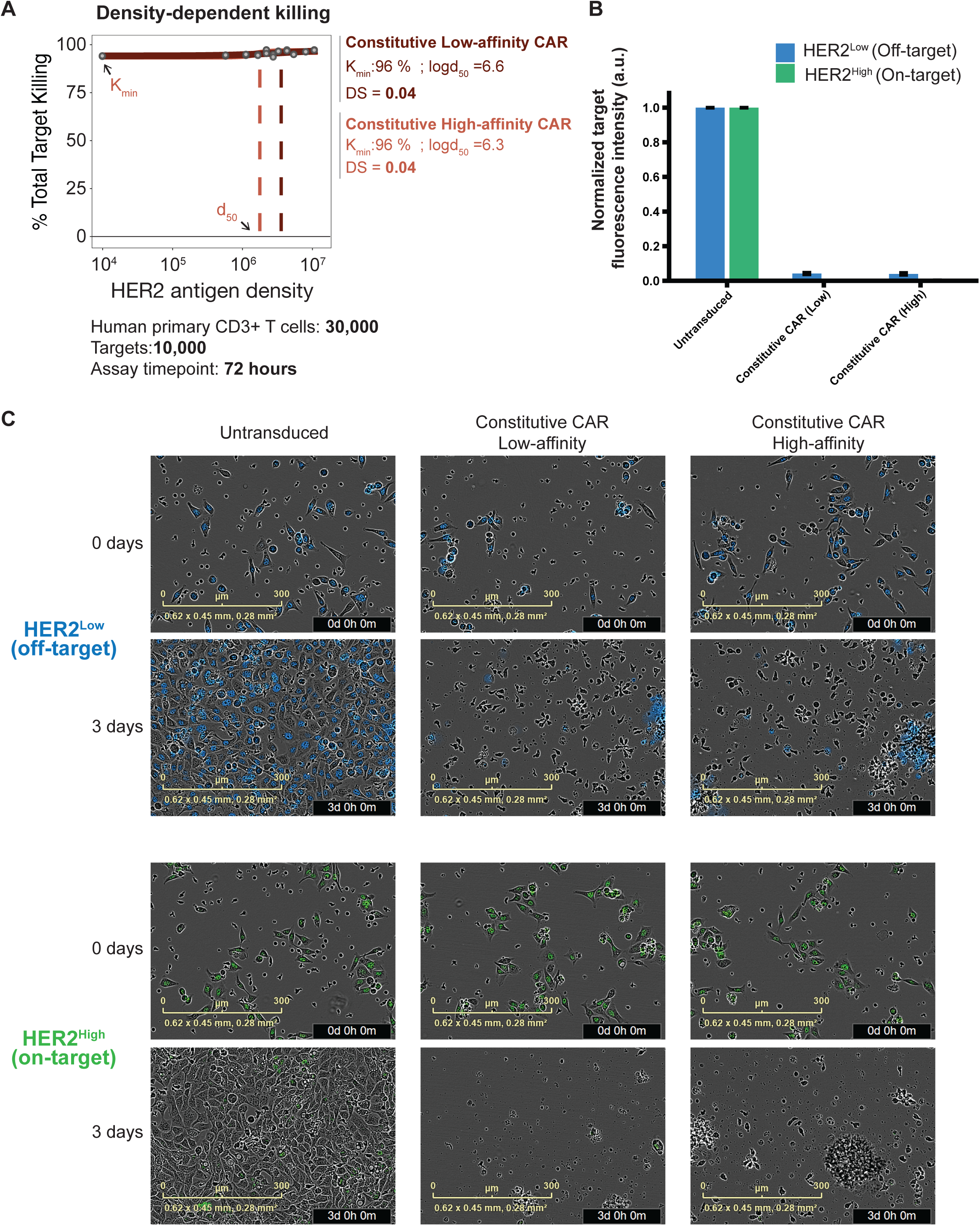
Representative timelapse imaging and analysis of killing activity for high- and low-affinity constitutive CARs. **(A)** *In vitro* Total target killing comparing constitutive anti-HER2 CAR of low- and high-affinity as a function of target antigen density. **(B)** Normalized target fluorescent intensity in co-culture assays of high-antigen (on-target) or low-antigen (off-target) HER2 targets with untransduced CD3+ T cells, or engineered T cells with constitutive high-affinity or low-affinity anti-HER2 CAR after 72 hours. bars represent mean ± SEM (n =3 replicates). (**C**) Representative images at the beginning (t=0 days) and end (t = 3 days) of co-culture killing assays.

**Figure S3:**
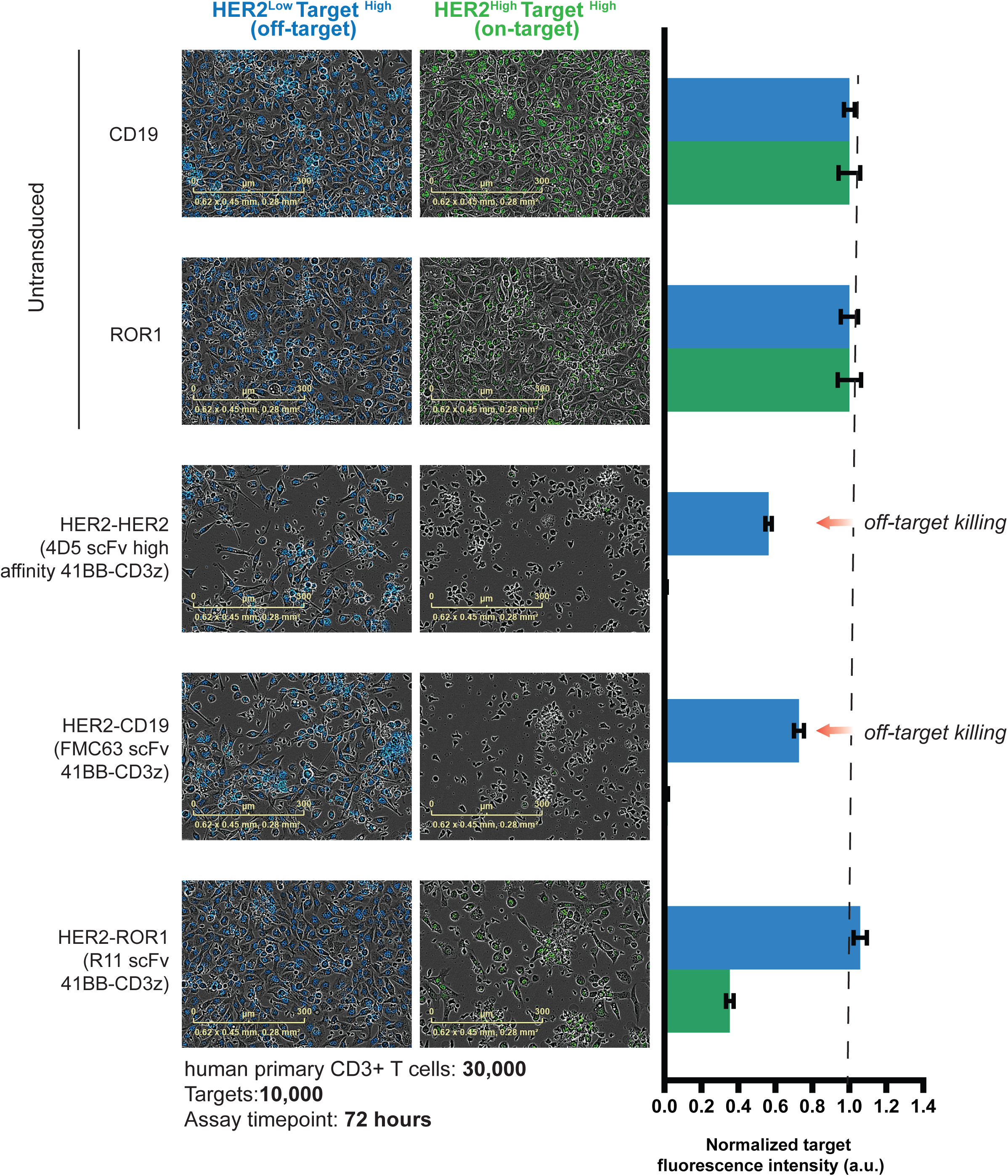
Representative timelapse imaging and analysis of killing activity for SynNotch CAR circuits. Representative microscopy images at the end of co-culture killing assays (t= 3 days) comparing untransduced T cells and inducible SynNotch-CAR circuits against their respective off-target or on-target cell lines. Right: normalized target fluorescent intensity for on-target and off-target cells. Bars represent mean ± SEM (n =3 replicates).

**Figure S4:**
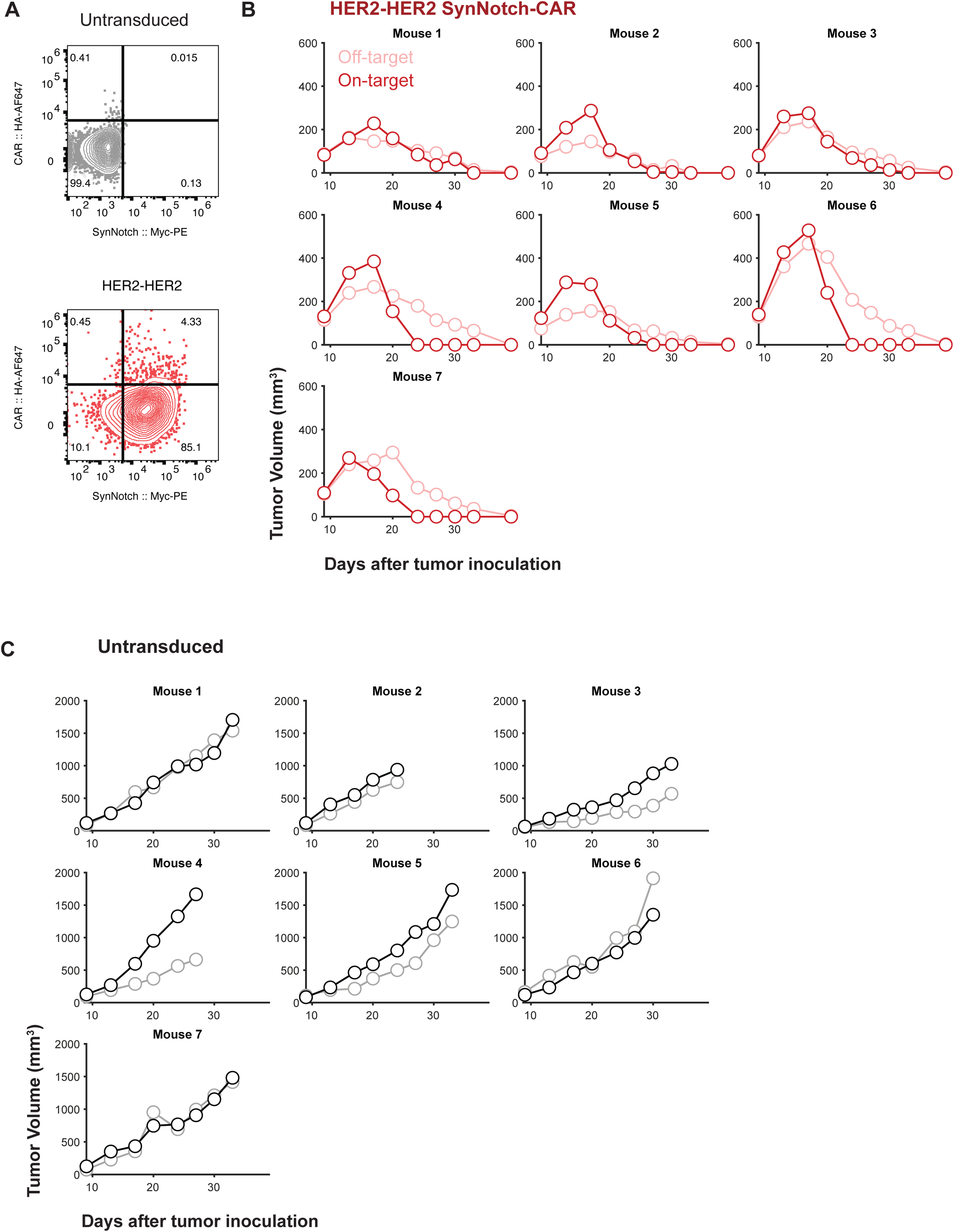
Individual mouse data, related to Figure 1G. **(A)** Representative Flow cytometry plots showing synNotch and basal CAR expression (%HA+) of T cells pre-injection into xenograft mouse model. **(B-C)** Individual tumor curves of mice bearing a HER2^Low^ tumor (off-target flank) and HER2^High^ tumor (on-target flank) and treated with 5M untransduced CD3+ T cells or WT HER2-HER2 synNotch-CAR T cells (n=7/group).

**Figure S5:**
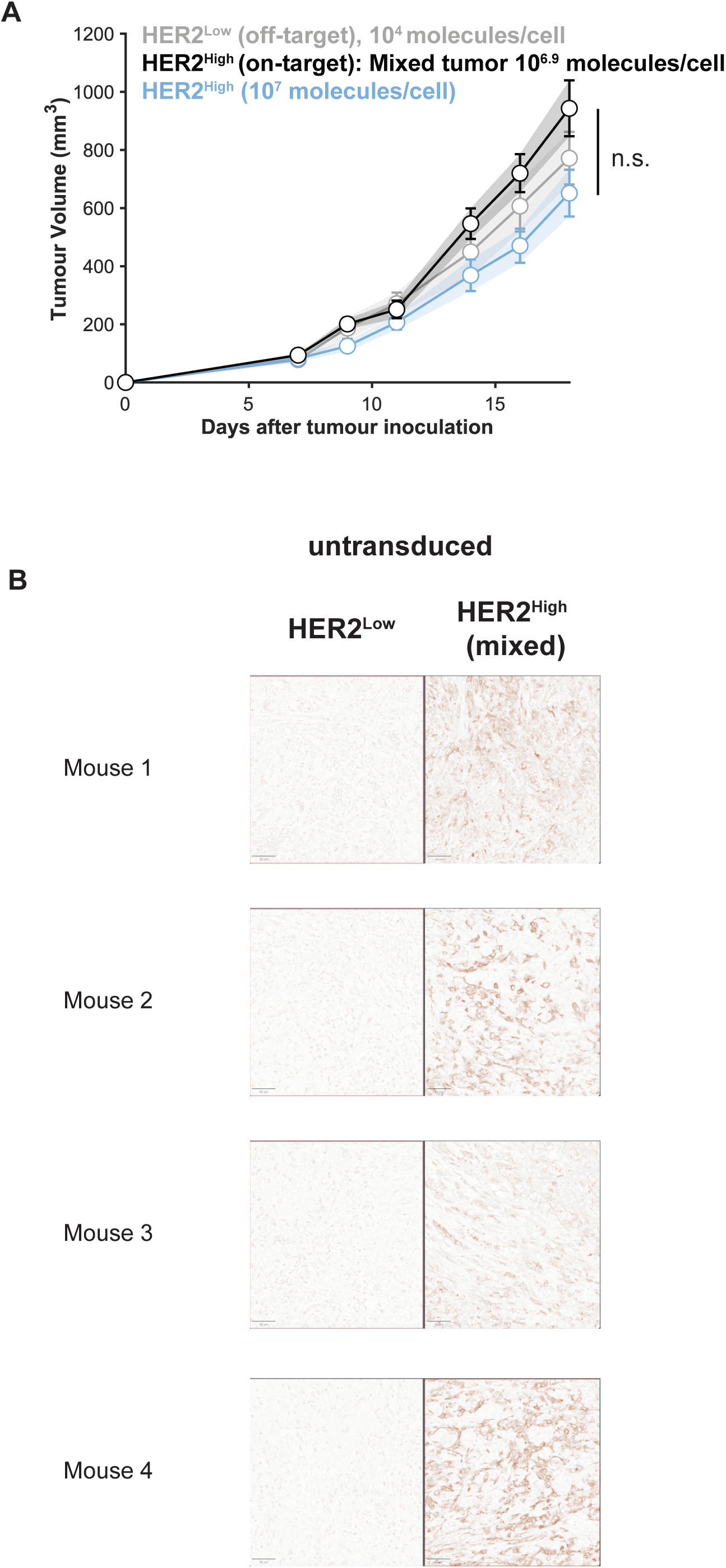
Tumor growth of single tumors, related to Fig 1G. **(A)** Tumor **g**rowth curves of either HER2^Low^, HER2^High^ or a mixed tumor (75% HER2^High^, 25% HER2^Low^) in mice treated with untransduced T cells. **(B)** Representative immunohistochemistry images of HER2 in tumors recovered at the end of the experiments from the untransduced group. The tumor composition did not change over time.

**Figure S6:**
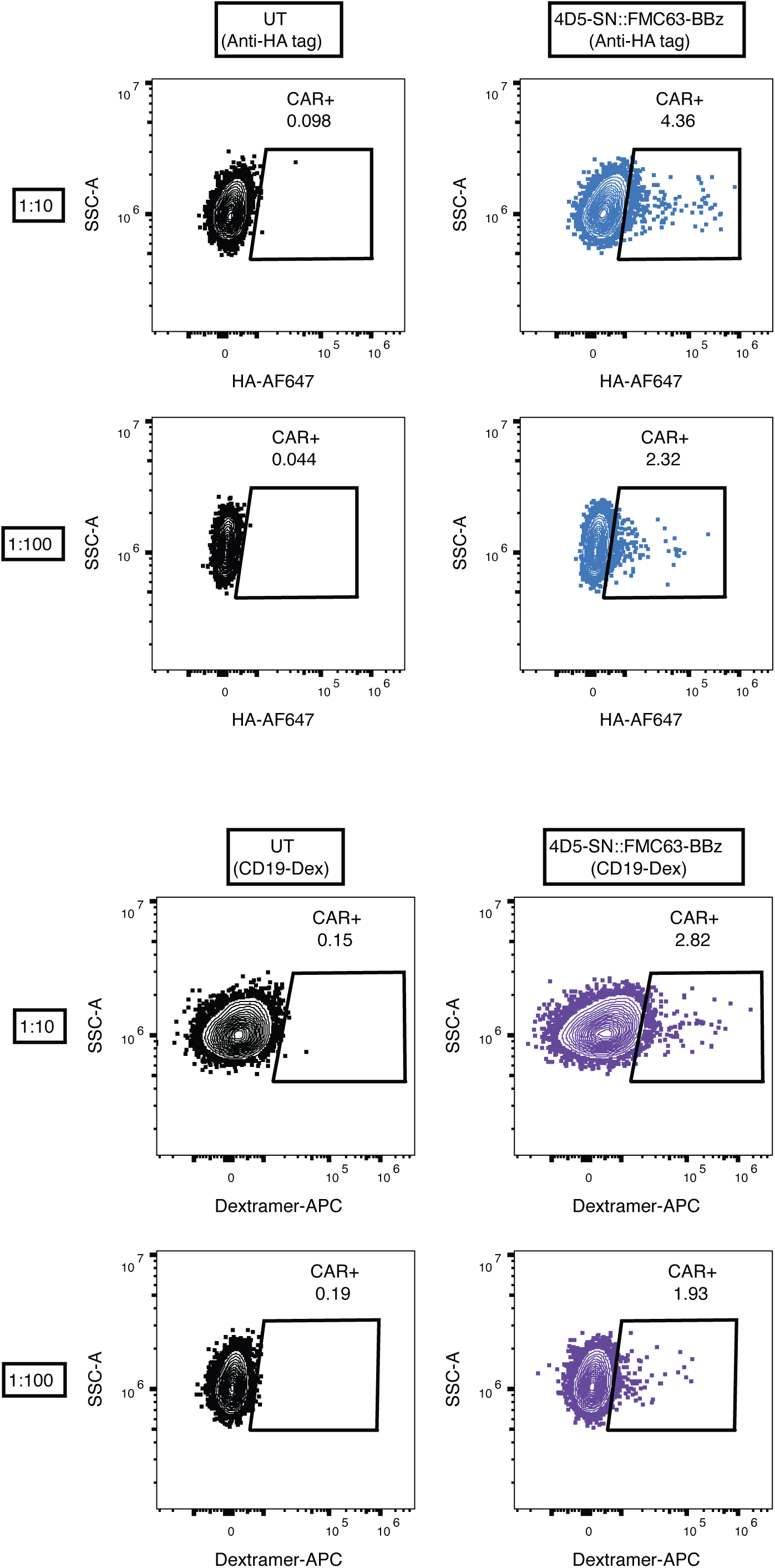
Comparison of epitope HA Tag and CD19-dextramer for measuring CAR surface expression, related to Fig. 1E. Representative flow plots comparing the basal CAR surface expression in HER2-CD19 SynNotch-to-CAR circuit and untransduced T cells using either an anti-HA antibody or a CD19-dextramer at two different staining dilutions.

**Figure S7.**
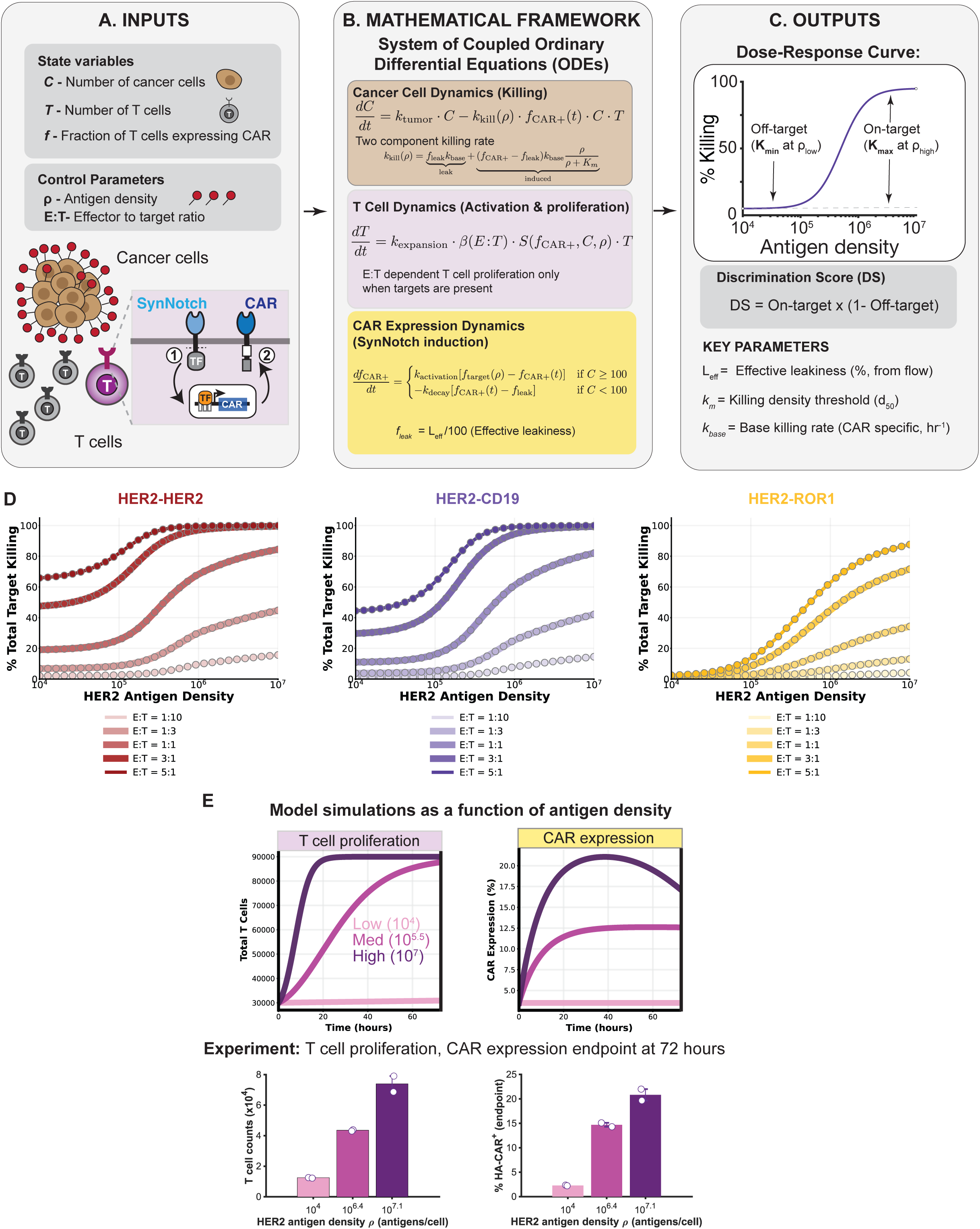
Details of mathematical model for synNotch-CAR T-cell killing. **(A–C)** Extended schematic diagram of the ordinary differential equation (ODE) modeling framework. **(A)** Model inputs including continuous state variables (*C*, tumor cancer cell count; *T*, total T-cell count; *f*, fraction of T cells expressing CAR) and control parameters (ρ, target antigen density in molecules/cell; E:T, Effector-to-Target cell ratio). Cellular schematic depicts target antigen binding to synNotch, induced transcription factor translocation, and subsequent CAR cell-surface expression. **(B)** System of coupled non-linear ODEs describing tumor cell killing dynamics (*dC/dt*), T-cell activation, E:T ratio-dependent logistic proliferation (*dT/dt*), and synNotch-driven CAR expression induction and decay kinetics (*df_CAR+/dt*). **(C)** Model outputs include simulated 4-parameter dose response curves highlighting off-target baseline killing (K_min_ at ρ_low_ = 10^4^) and on-target maximal killing (K_max_ at ρ_high_ = 10^7^), calculation of the Discrimination Score (DS = On-target * (1 − Off-target)), and summary of key kinetic parameters (*L_eff*, effective leakiness %; *K_m_*, density threshold *d_50_*; k_base_, baseline CAR killing rate). **(D)** Simulated 72-hour dose-response killing curves across a 50-fold sweep of initial Effector-to-Target ratios (E:T = 1:10, 1:3, 1:1, 3:1, 5:1) for HER2-HER2, HER2-CD19, and HER2-ROR1 synNotch-CAR circuits. **(E)** Comparison of model time-course simulations (top panels) and 72-hour experimental endpoint measurements (bottom panels) for the HER2-CD19 circuit across Low (10^4^), Med (10^5.5^ or 10^6.4^), and High (10^7^ or 10^7.1^) HER2 antigen densities. Top plots show simulated T-cell proliferation and CAR expression percentage over 72 hours. Bottom bar plots display experimental T-cell counts and percentage HA-CAR+ cells measured at 72 hours post-co-culture. Circles represent individual experimental replicates (n=2).

**Figure S8.**
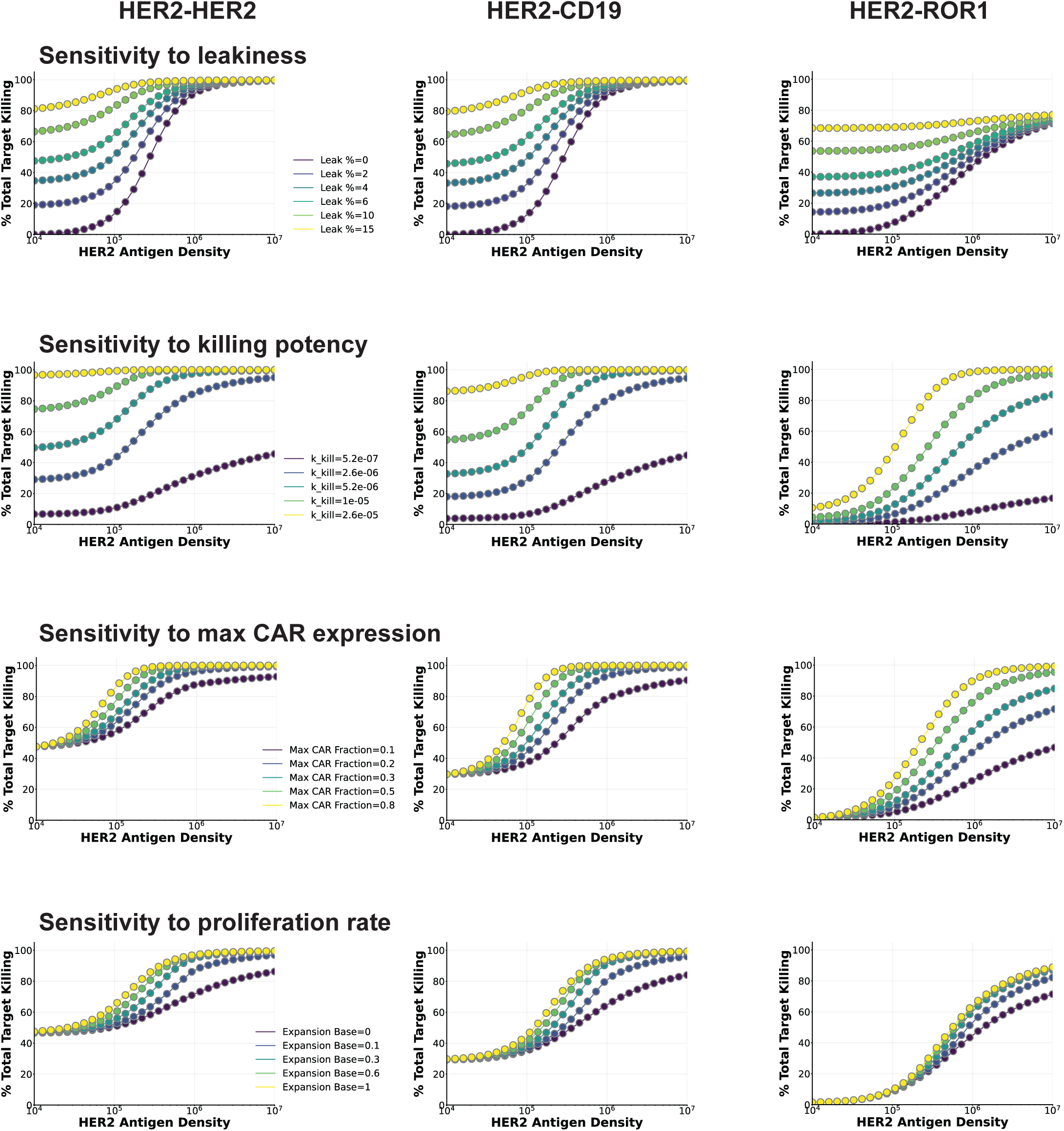
Sensitivity analysis for model parameters of SynNotch-CAR T cell function, related to Fig. 2E. Systematic single-parameter sensitivity sweeps evaluating the effect of core model parameters on 72-hour dose-response target killing curves across three calibrated synNotch-CAR constructs: HER2-HER2, HER2-CD19, and HER2-ROR1. **Row 1 (Sensitivity to leakiness):** Impact of varying baseline uninduced leak fraction (Leak % = 0%, 2%, 4%, 6%, 10%, 15%). **Row 2 (Sensitivity to killing potency):** Impact of varying base single-cell cytotoxicity constant k_kill,base (5.2×10^-7 to 2.6×10^-5 cell^-1 h^-1). **Row 3 (Sensitivity to max CAR expression):** Impact of varying maximal CAR fraction ceiling (f_max = 0.1, 0.2, 0.3, 0.5, 0.8). **Row 4 (Sensitivity to proliferation rate):** Impact of varying baseline antigen-driven expansion rate (α_base = 0.0, 0.1, 0.3, 0.6, 1.0).

**Figure S9.**
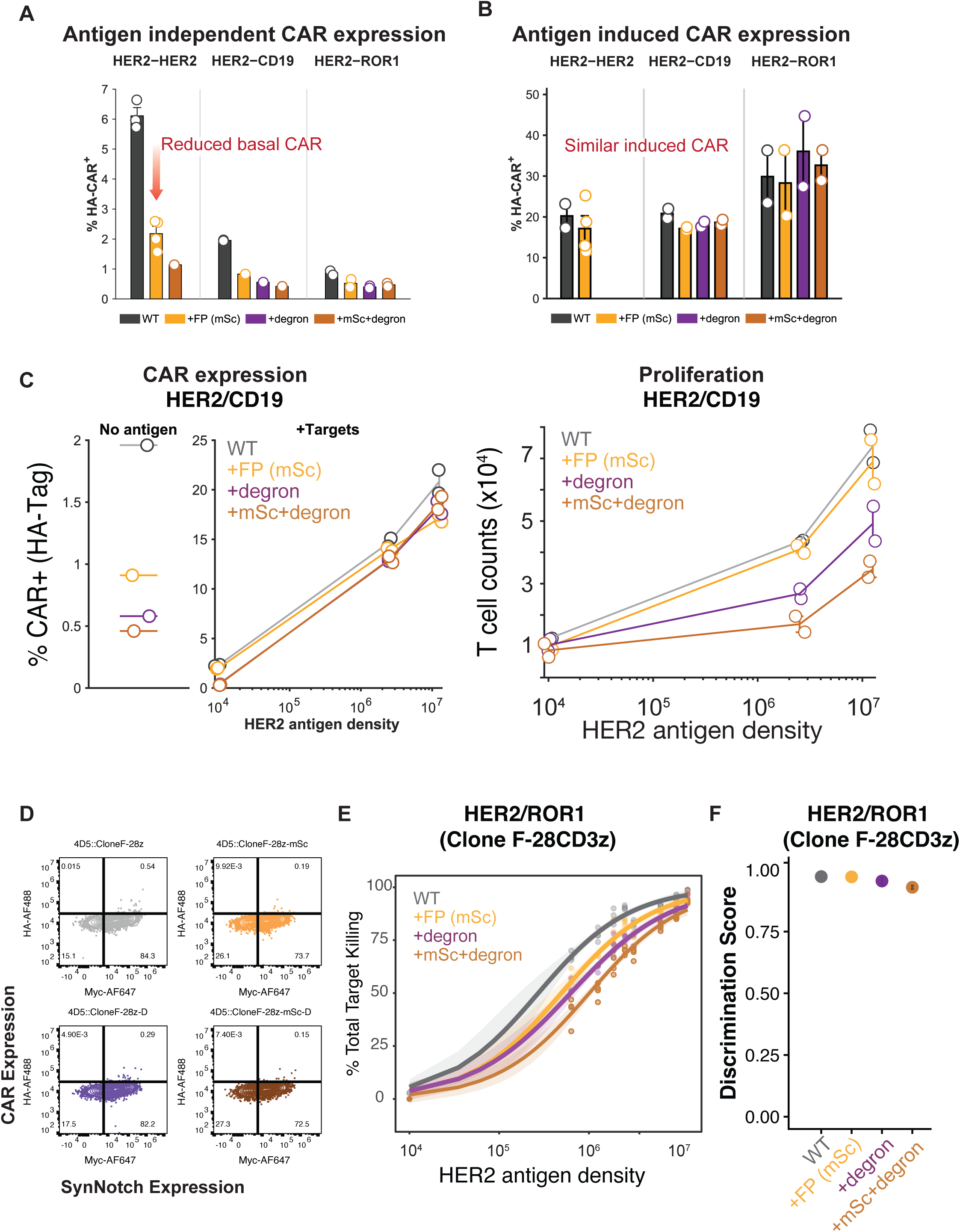
CAR expression, proliferation and discrimination score, related to Fig 3. **(A)** Flow cytometry quantification of antigen-independent (leaky) CAR expression percentage (% HA-CAR+) measured 10 days post CD3/CD28 bead activation for HER2-HER2, HER2-CD19, and HER2-ROR1 SynNotch-CAR circuits. Circles represent individual experimental replicates; Bars represent mean ± SEM. **(B)** Flow cytometry quantification of antigen-induced CAR expression percentage (% HA+) measured 72 hours post-induction with high-antigen density targets (10^7^ HER2 molecules/cell) across one-layer WT circuit and two-layer circuit variants. All variants maintain similar levels of maximal induced CAR expression. Circles represent individual experimental replicates; Bars represent mean ± SEM. **(C)** Experimental characterization of HER2-CD19 circuit modifications across targets with three different antigen densities (10^4^ to 10^7^ molecules/cell). Left panel depicts CAR expression percentage (% CAR+ HA-Tag) in the absence of antigen (leaky CAR) and upon target antigen exposure (+Targets). Right panel displays T cell counts after 72 hours of co-culture (T cell counts x 10^4^) with targets expressing different antigen densities. **(D)** Representative flow cytometry plots for synNotch and basal CAR expression for HER2-ROR1 one-layer and two-layer circuits. **(E)** Density dependent total killing for synNotch-CAR HER2-ROR1 circuit variants, each dot represents an individual well, curves were fitted to the data using a 4-parameter logistic curve and shaded areas show confidence bands. **(F)** Discrimination Score (DS) for HER2-ROR1 one-layer WT circuit (gray) and two-layer circuit variants. The ROR1 CAR has a binding domain derived from clone F and a CD28 transmembrane domain and co-stimulation domain with a CD3ζ signaling domain.

**Figure S10.**
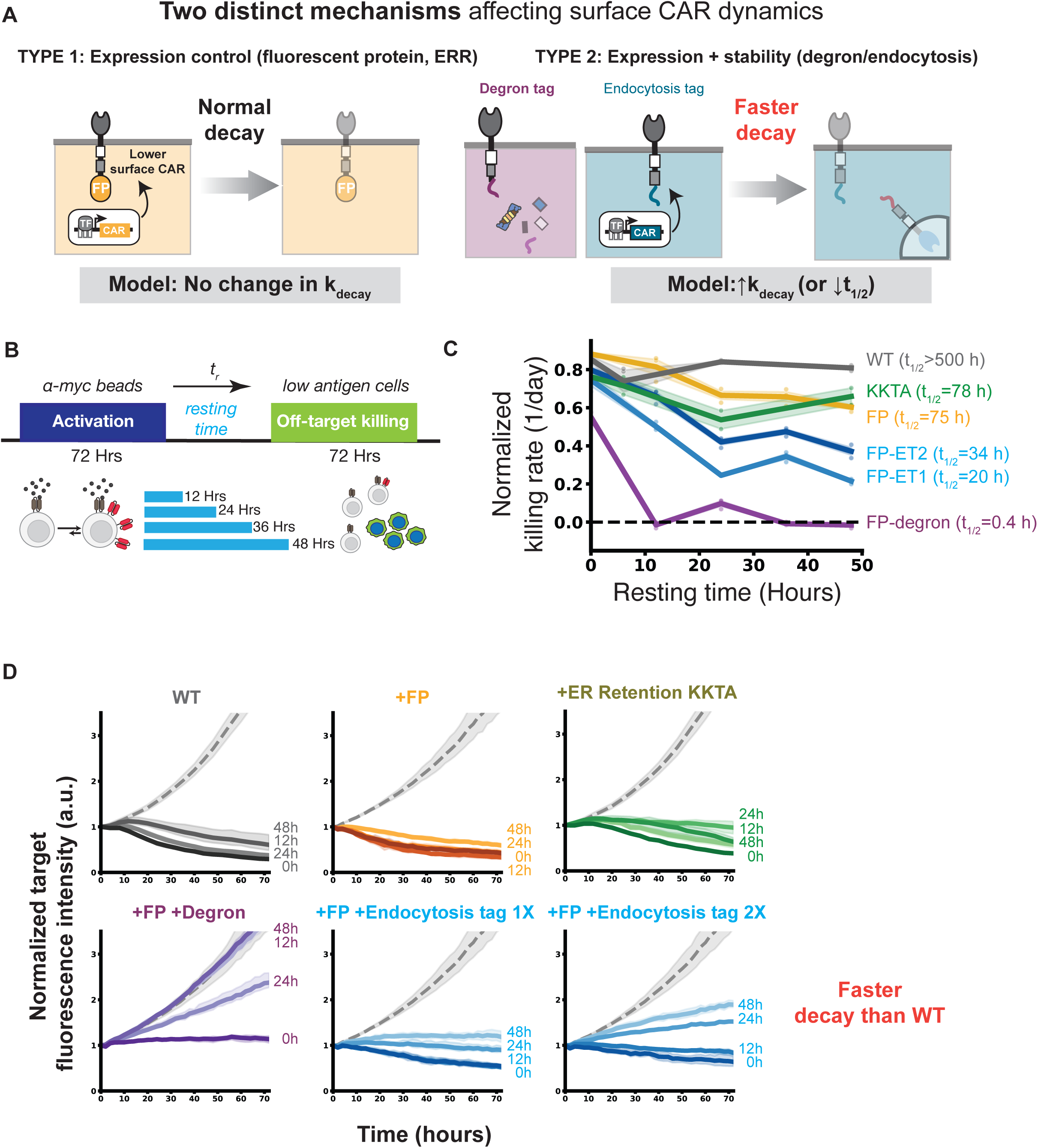
Post-translational modules tune synNotch-induced CAR surface expression and killing half-life. **(A)** Schematic depicting two distinct post-translational mechanisms modulating cell-surface CAR dynamics. Type 1 (Expression control, Fluorescent Protein tag and ER retention tag): lower basal CAR expression and insertion into the plasma membrane with degradation kinetics (k_decay_) and killing activity upon induction similar to WT. Type 2 (Expression + stability control, Degron or Endocytosis tags): Accelerated CAR degradation and endocytic internalization leading to increased degradation rate constant (faster k_decay_) and decreased surface half-life (t_1/2_). **(B)** Schematic of the activation, resting-time and cytotoxicity assay workflow. synNotch-CAR T cells were activated with anti-myc beads for 72 hours to induce CAR expression, beads were removed and rested in media for variable time intervals (t_r_ = 0, 12, 24, 36, 48 hours), and subsequently co-cultured with low-antigen off-target cells in a 72-hour cytotoxicity assay. **(C)** Normalized off-target killing rate (1/day) as a function of resting time (hours) across one-layer WT circuit and two-layer circuit variants exhibiting distinct CAR surface half-lives (t_1/2_). **(D)** Time course of normalized target fluorescence intensity (a.u.) over 70 hours co-culture assay with low-antigen off-target cells following 0h, 12h, 24h, and 48h resting periods. Profiles are shown for one-layer, WT and two-layer HER2-HER2 circuit variants.

**Figure S11:**
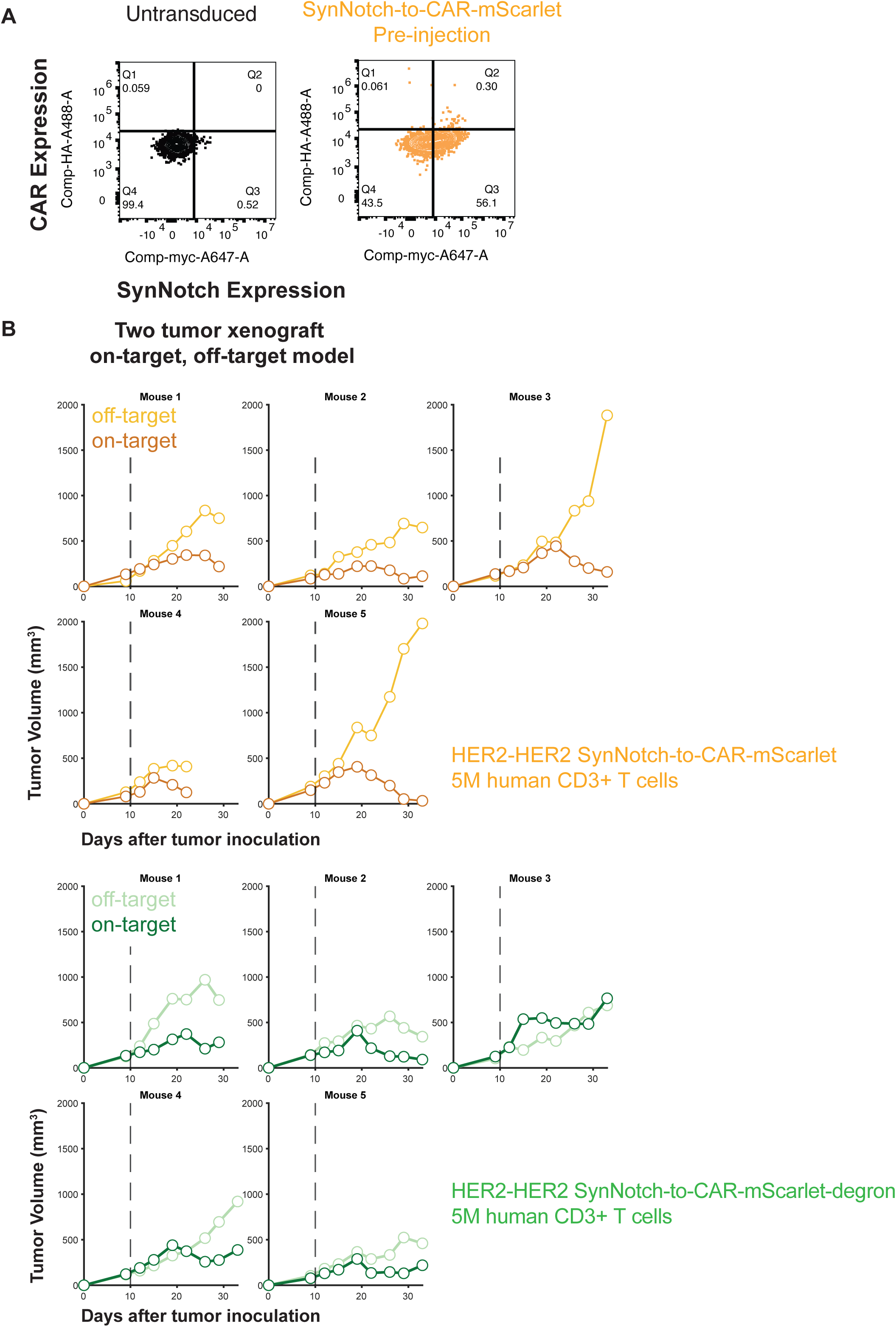
SynNotch expression, basal CAR expression and individual mouse data, related to Fig. 4A. **(A)** Preinjection receptor expression for synNotch and basal CAR expression of control CD3+ T cells and HER2-HER2+mScarlet T cells as measured by flow cytometry. **(B)** Individual tumor curves of dual-xenograft bearing mice (left: HER2^low^ off-target flank, right: HER2^high^ on-target flank) receiving 5 million HER2-HER2+mSc T cells or HER2-HER2+mSc+degron T cells (n=5/group).

**Figure S12:**
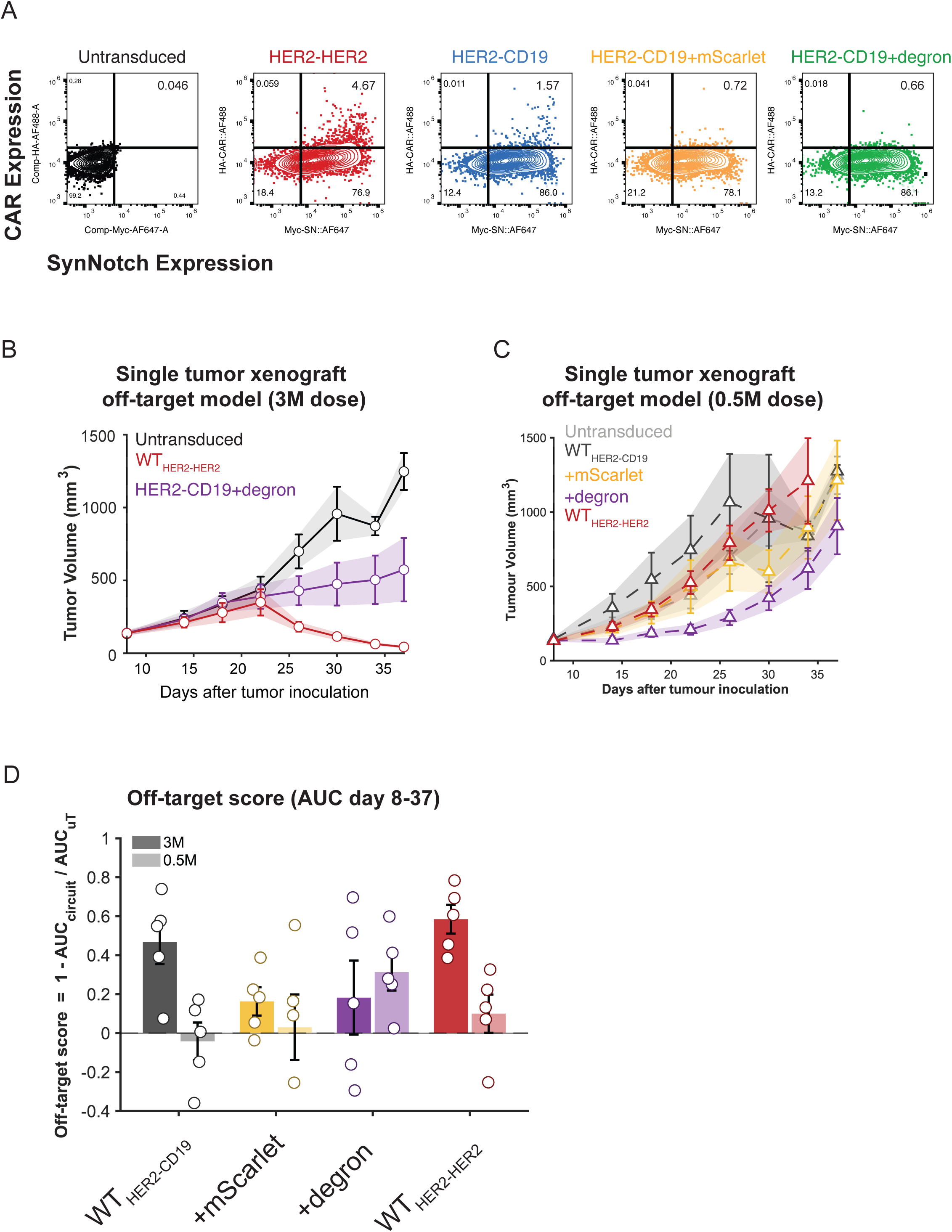
SynNotch expression, basal CAR expression and off-target score, related to Fig. 4C-D. **(A)** Preinjection receptor expression for synNotch and basal CAR expression of control CD3+ T cells, WT HER2-HER2 T cells, WT HER2-CD19 T cells, HER2-CD19+mSc T cells, and HER2-CD19+degron T cells as measured by flow cytometry. **(B-C)** Tumor volumes of mice bearing a single HER2^low^ CD19^high^ tumor (off-target model) and treated with control CD3+ T cells, WT HER2-HER2 T cells, WT HER2-CD19 T cells, HER2-CD19+mSc T cells, or HER2-CD19+degron T cells (n=5/group) at the **(B)** 3 million dose or **(C)** 0.5 million dose. **(D)** Off-target score calculated from the tumor volume area under curve from Day 8 to Day 37 of various treatment groups: WT HER2-HER2 T cells, WT HER2-CD19 T cells, HER2-CD19+mSc T cells, or HER2-CD19+degron T cells (n=5/group) at the 3 million dose or 0.5 million dose. Individual data points represent single mice; bars represent mean ± SEM.

**Figure S13:**
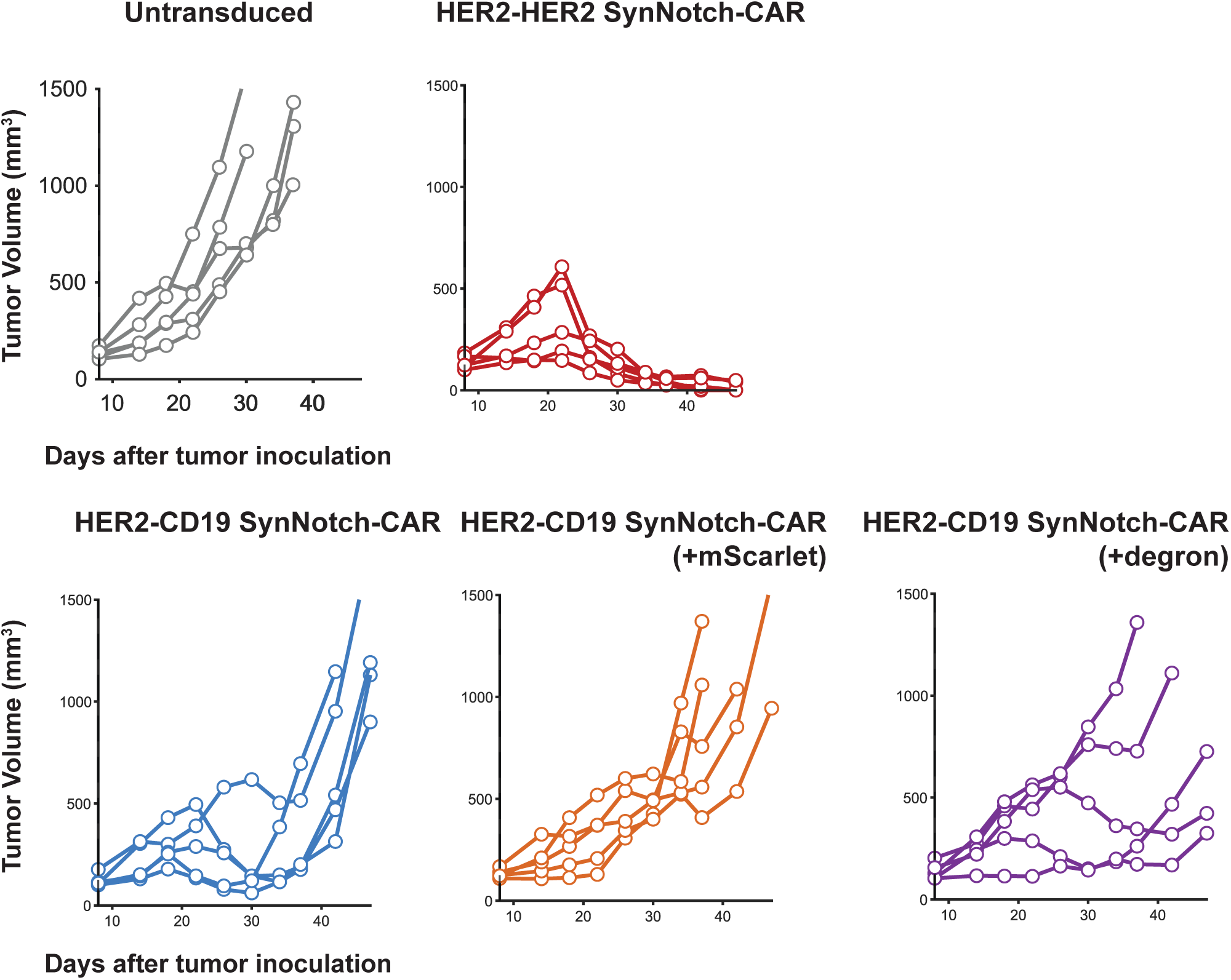
Individual mouse data, related to Fig. 4C-D. Individual tumor curves of mice bearing a single HER2^low^ CD19^high^ tumor (off-target model) and treated with 3M control CD3+ T cells, WT HER2-HER2 T cells, WT HER2-CD19 T cells, HER2-CD19+mScarlet T cells, or HER2-CD19+degron T cells (n=5/group).

**Figure S14:**
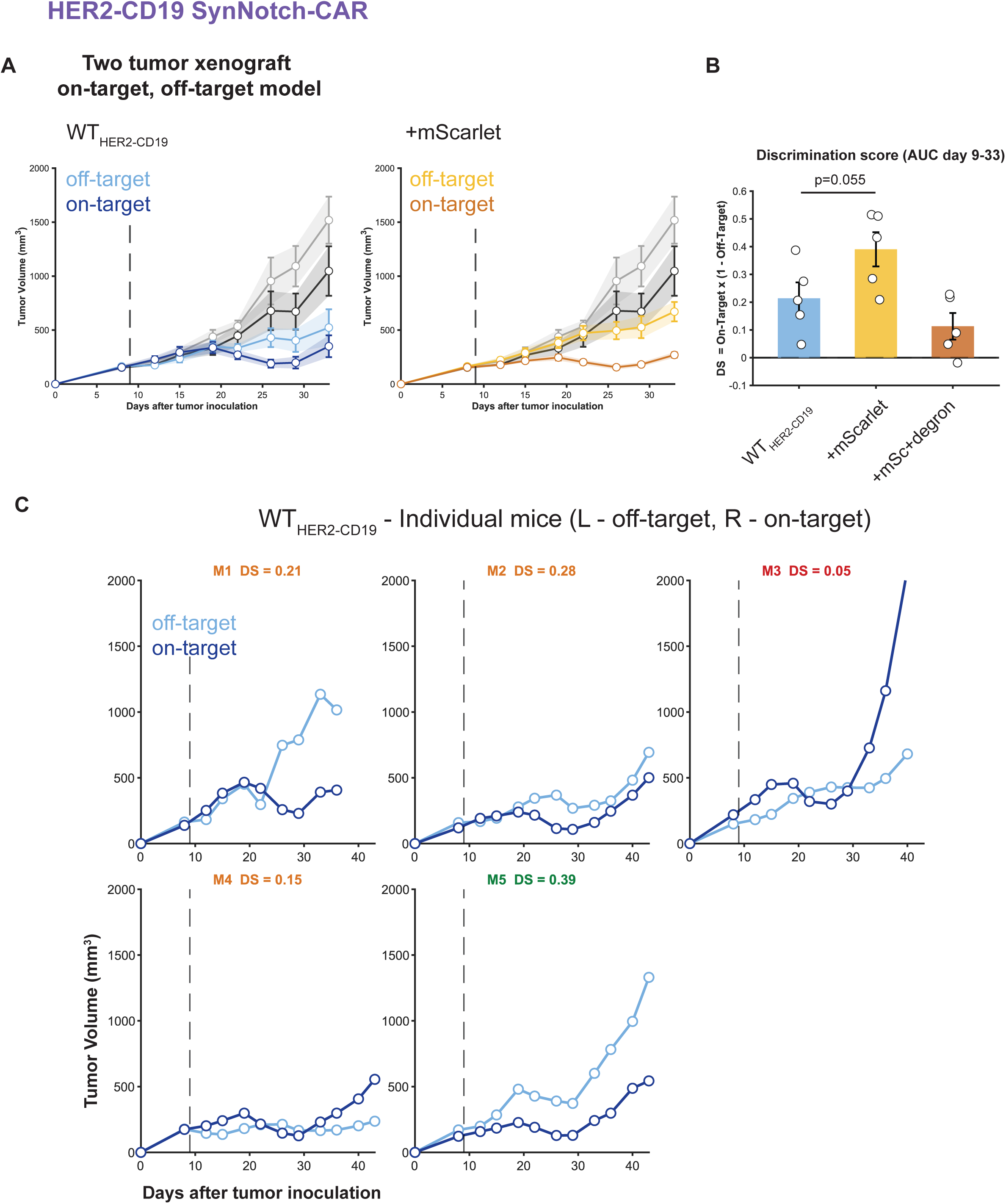
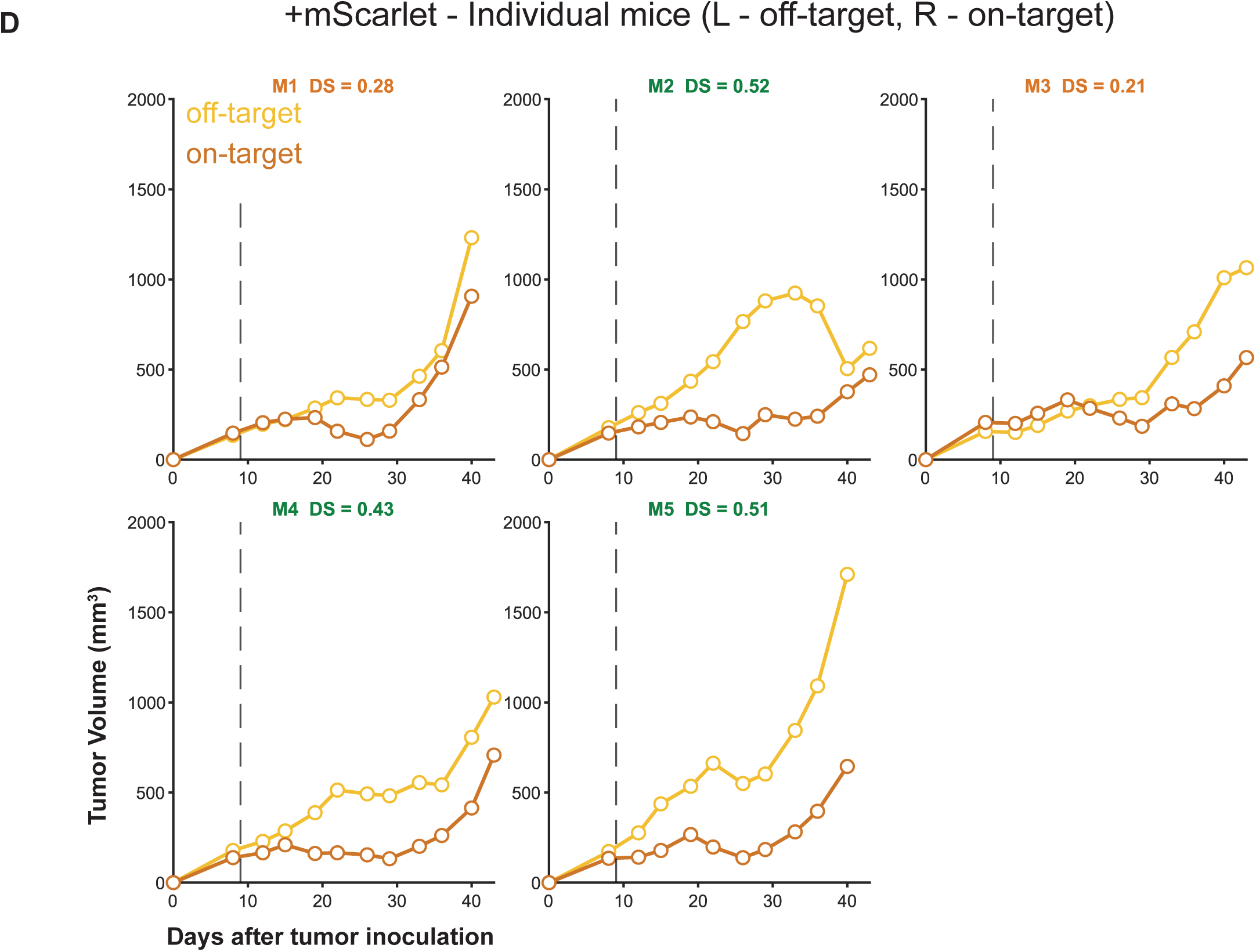
Additional two-tumor mouse model to evaluate HER2-CD19 SynNotch-CAR circuit, related to Fig. 4A. **(A)** Tumor curves of dual-xenograft bearing mice with 1 million of HER2^low^ CD19 (off-target, left flank) and 1 million of 25% HER2^low^ CD19/75% HER2^high^ CD19 mixture (on-target, right flank) and treated with 3 million of WT HER2-CD19 T cells or HER2-CD19+mSc T cells (n=5/group). **(B)** Discrimination score calculated based on tumor volume area under curve (AUC) from Day 9 to Day 33 of various treatment groups: 3M WT HER2-CD19 T cells, HER2-CD19+mSc T cells, or HER2-CD19+mSc+degron (n=5/group). Individual data points represent single mice; bars represent mean ± SEM. Group comparisons were performed using the Mann-Whitney U test. **(C)** Individual tumor curves of mice treated with 3 million WT HER2-CD19 T cells or **(D)** HER2-CD19+mSc T cells (n=5/group).

**Figure S15:**
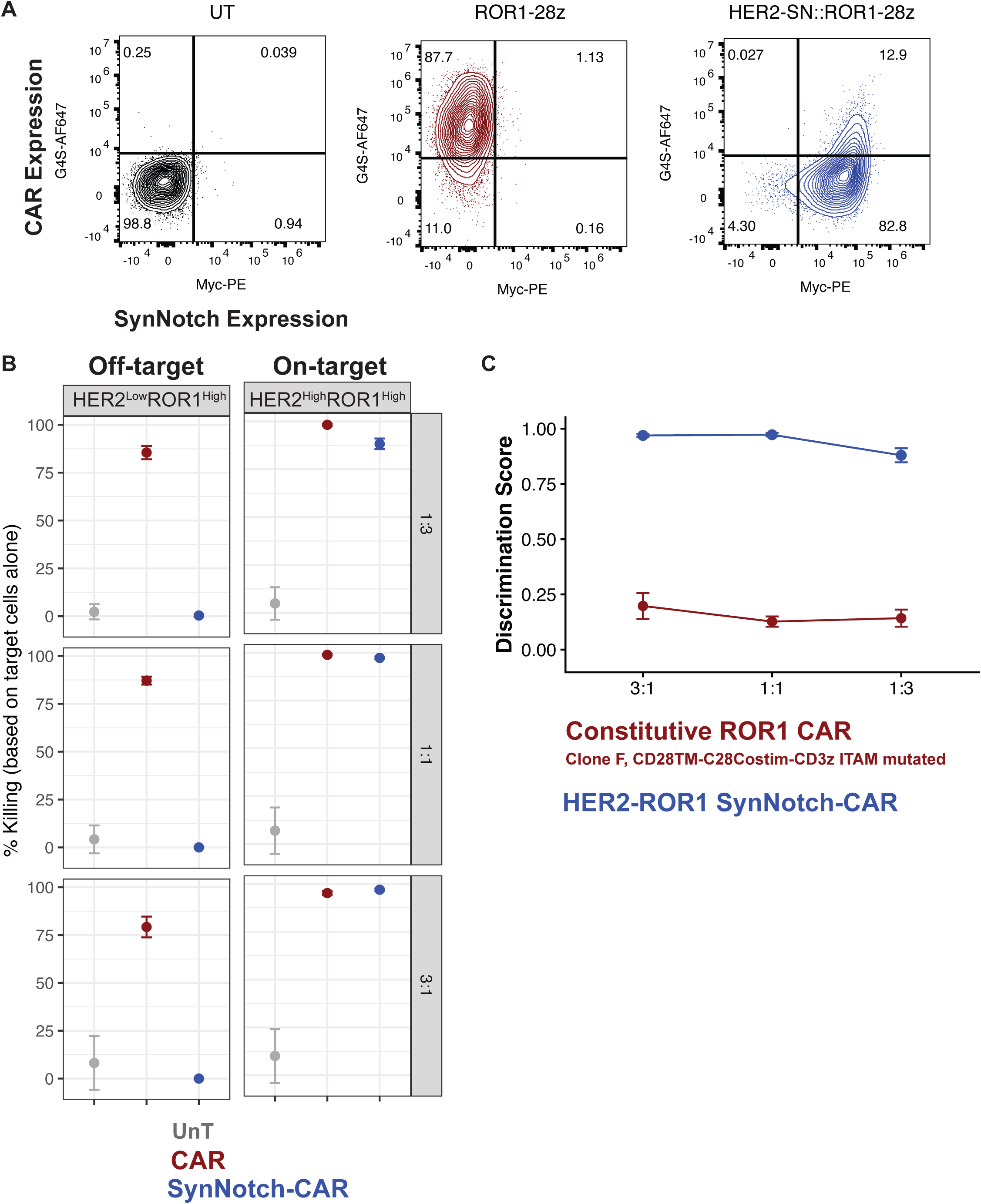
Additional data for HER2-ROR1 SynNotch-CAR circuit. **(A)** Receptor expression for synNotch and CAR expression of control CD3+ T cells, constitutive ROR-28z CAR T cells, and inducible HER2-ROR1-28z synNotch-CAR T cells as measured by flow cytometry **(B)** Percent specific lysis of cancer cell lines (HER2^low^ ROR1^high^ HCC1806 off-target cancer cells or HER2^high^ ROR1^high^ HCC1806 on-target cancer cells) when co-cultured with control CD3+ T cells, constitutive ROR-28z CAR-T cells, and inducible HER2-ROR1-28z synNotch-CAR T cells at an E:T (Effector-to-Target) ratio of 1:3, 1:1, and 3:1. **(C)** Discrimination score (DS) calculated based on the specific killing of HER2^high^ ROR1^high^ HCC1806 (on-target) and HER2^low^ ROR1^high^ HCC1806 (off-target) cancer cells. In vitro assays are performed in three technical replicates. Data are shown as mean ± SEM (n=3).

**Figure S16:**
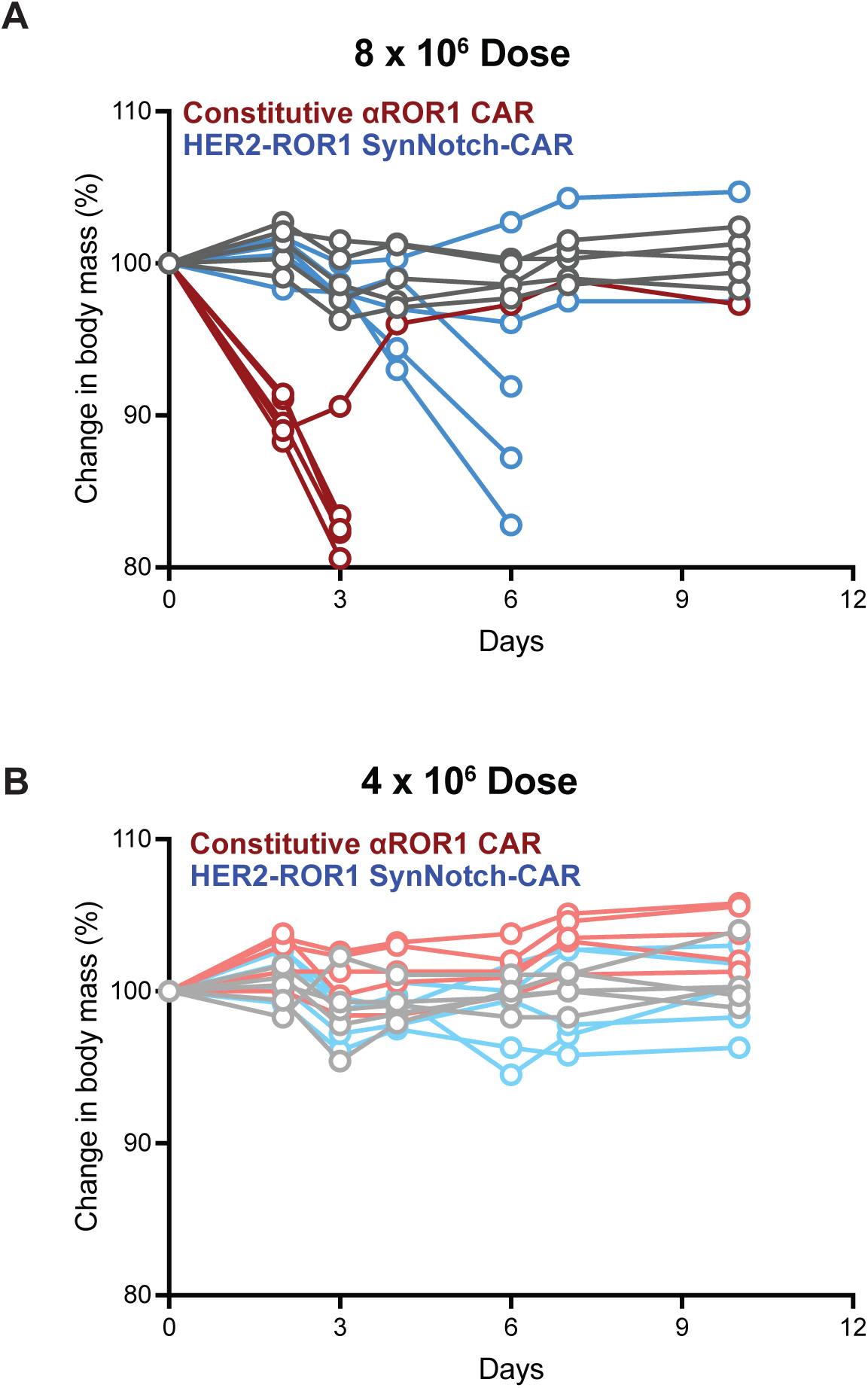
Toxicity model to evaluate HER2-ROR1 SynNotch-CAR circuit. **(A-B)** Change in body mass of tumor-free mice treated with control CD3+ T cells, constitutive ROR-28z CAR-T cells, or HER2-ROR1-28z synNotch-CAR T cells at the **(A)** 8 million dose or **(B)** 4 million dose (n=5/group).

**Figure S17:**
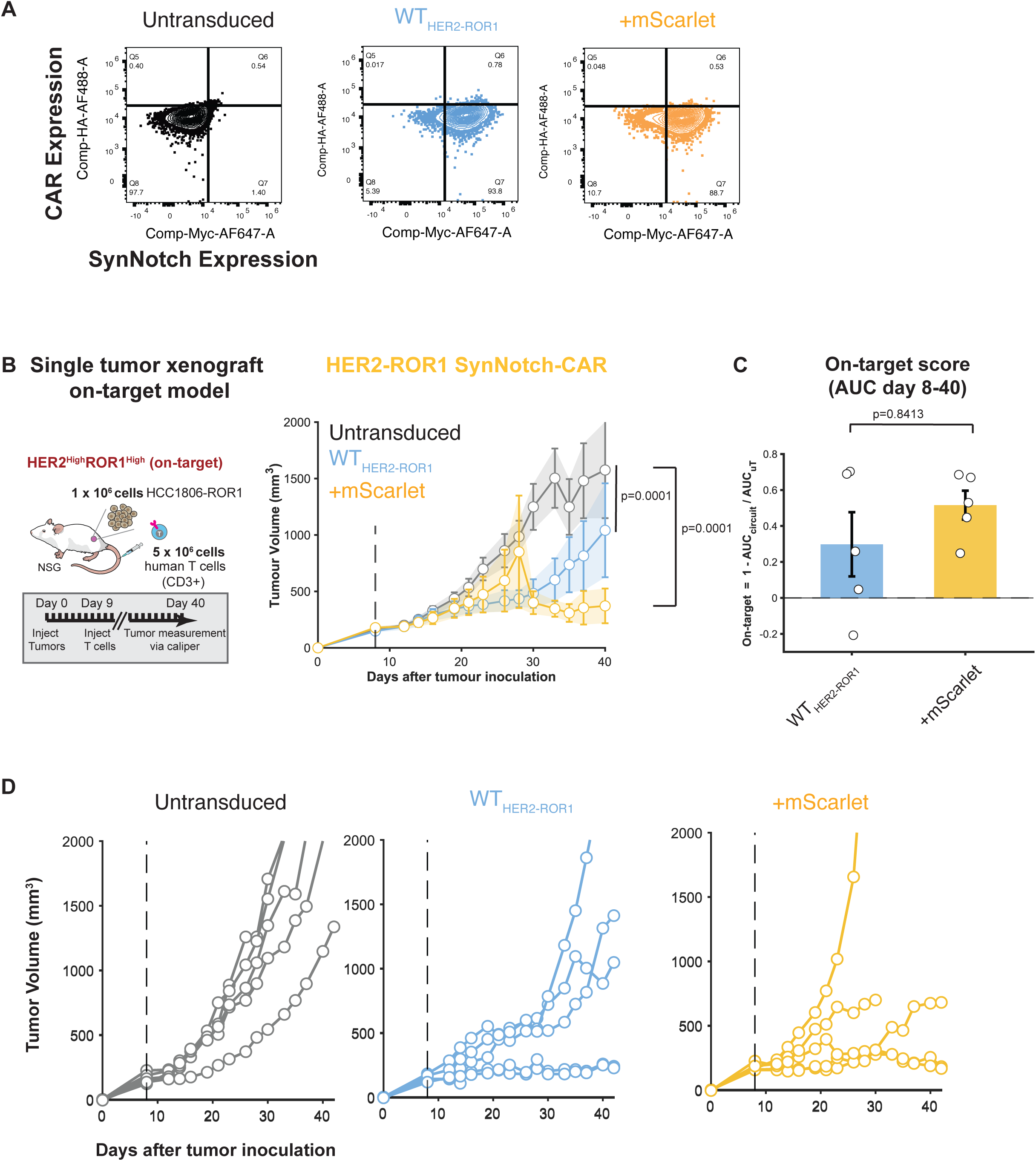
Additional single-tumor mouse model HER2-ROR1 SynNotch-CAR circuit. **(A)** Preinjection receptor expression for synNotch and basal CAR expression of control CD3+ T cells, WT HER2-ROR1 T cells, and HER2-ROR1+mScarlet T cells as measured by flow cytometry. **(B)** Schematic of experiments performed with mice bearing a single engineered human breast cancer xenograft of 1 million HER2^low^ ROR1^high^ HCC1806 (on-target model). Tumor volumes of mice treated with 5 million untransduced CD3+ T cells, WT HER2-ROR1 T cells, or HER2-ROR1+mSc T cells (n=5/group). **(C)** On-target score calculated based on the tumor volume area under curve (AUC) from Day 8 to Day 40 of mice receiving 5 million WT HER2-ROR1 T cells or HER2-ROR1+mSc T cells (n=5/group). Individual data points represent single mice; bars represent mean ± SEM. Group comparisons were performed using the Mann-Whitney U test. **(D)** Individual tumors curves for mice receiving 5 million control T cells, WT HER2-ROR1 T cells, or HER2-ROR1+mSc T cells (n=5/group).

