## Supplemental Material for "Rational Control of Basal CAR Expression Improves Discrimination in Inducible T Cell Circuits"

**MATERIAL AND METHODS**

**Gene synthesis and cloning**

Gene fragments were codon optimized using the tool on the Integrated DNA Technologies (IDT) website and obtained as gene fragments from Twist Bioscience. All lentiviral constructs were synthesized using in-fusion cloning (Takara Bio #638943) and Stellar Competent Cells (Takara Bio #636766).

*Constitutive CAR and synNotch-CAR circuit designs*

Anti-HER2 SynNotch and CAR constructs were built using scFvs derived from clone 4D5^1^, anti-CD19 SynNotch and CAR constructs were built using the scFv derived from FMC63, and anti-ROR1 CAR constructs were built using scFvs derived from clone R11^3^ and clone F^2,3^ .

Anti-HER2 and anti-CD19 CARs were synthesized by fusing the N-terminal CD8α signaling peptide (MALPVTALLLPLALLLHAARP) and an HA-tag (YPYDVPDY) to the scFv binder, hinge region of the human CD8α chain, transmembrane and intracellular regions of the human 4-1BB costimulatory domain, and human CD3ζ signaling domains.

For the ROR1 CAR variants shown in Figure 1 and S3, we used the clone R11 ScFv, CD8 transmembrane, 4-1BB costimulatory domain and the human CD3ζ signaling domain. For the variants in Figure S9, we used the Clone F scFv, CD28 transmembrane and CD28 costimulatory domain together with the human CD3ζ signaling domain. For Figures S15 and S16, we used the ROR1 CAR sequence previously reported by the Mackall Lab which has a GM-CSF signaling peptide, clone F scFv, CD28 transmembrane domain, CD28 costimulatory domain and a modified human CD3ζ signaling domain^2^. For the experiment shown in Figure S17, the CAR sequence was similar to the one used in Fig. S16 but incorporated an HA-Tag.

**C-terminal tags:** The degron sequence used in this study was obtained from the C-terminal region of the mouse ornithine decarboxylase (cODC) domain and previously used in a CAR as a fusion^1^. The endocytosis sequence was obtained from the C-terminal region of the Cytotoxic T-Lymphocyte Associated Protein 4 (CTLA-4) tail. ER retention motifs were RRR, KKYL, RKR, and KKTA. All C-terminal tags and fluorescent proteins (mCherry or mScarlet3) were fused to the CD3ζ signaling domain via a (GS)_4_ linker.

SynNotch receptors were synthesized by fusing the N-terminal CD8α signaling peptide (MALPVTALLLPLALLLHAARP) and myc-tag (EQKLISEEDL) with the scFv binder, mouse Notch1 minimal regulatory region, Gal4 DNA binding domain, and VP64 activation domain.

SNIPR receptors were synthesized by replacing the SynNotch Notch1 core regulatory domains with the SNIPR core, as described in Zhu et al. 2022^4^. The inducible promoter was also swapped from the miniCMV to the YB_TATA sequence (TCTAGAGGGTATATAATGGGGGCCA). The anti-CD19 binder for the SNIPR was the same clone FMC63 derived binder used for SynNotch.

SynNotch-CAR circuits were cloned into a single modified pHR’SIN:CSW vector to control synNotch to CAR copy number ratio at 1:1. SynNotch receptor expression is controlled by the pGK promoter, and CAR expression is controlled by a response element containing a 5-copy Gal4 DNA binding domain target sequence (GGAGCACTGTCCTCCGAACG) under the minimal CMV promoter as previously described^1^.

**Cancer cell line culture**

Engineered and parental HCC1806 (ATCC #CRL-2335) cell lines were cultured in Gibco^TM^ RPMI 1640 medium (ThermoFisher Scientific #11875119) and supplemented with 10% FBS (Cytiva #SH30071.03IH2540) until confluency before passage. Cells were washed with PBS and detached with TrypLE Express (ThermoFisher Scientific #12604021) via incubation at 37°C for 10-15 minutes. Fresh media was added to quench the TrypLE Express, and cells were centrifuged at 400g for 4 minutes, resuspended in fresh media, and replated in new flasks with more fresh media. Cell lines were routinely tested for mycoplasma contamination and maintained for a limited number of passages after thawing.

**Engineered HCC1806 breast cancer cell lines**

Engineered cell lines were derived from the HCC1806 breast cancer cell line (ATCC #CRL-2335). HCC1806 cells were transduced with lentivirus to express a SFFV promoter-driven nuclear fluorescent protein (miRFP or mEmerald) fused to the histone H2B for live-cell imaging. Additional target antigens are overexpressed with separate lentiviral vectors under SFFV promoters, including the extracellular and transmembrane region of human HER2 (AA 23-675) fused to tagBFP (Addgene Plasmid #164823), the extracellular region of human CD19 (AA 20-291) fused to the human CD19 transmembrane and intracellular domain (AA 301-556), and the extracellular region of human ROR1 (AA 30-406) fused to the human CD19 transmembrane domain (AA 301-556). Cells were sorted on a BD FACSAria II (BD Biosciences) based on fluorescence and surface antigen expression, expanded, and cryopreserved in 50% RPMI, 40% FBS, and 10% DMSO.

**Antigen abundance quantification**

Average antigen density per target cell was determined by quantitative flow cytometry, as previously described^1^. Briefly, 1 x 10^5^ cells from each target-cell population were stained with antigen-specific antibodies for 30 min on ice in 96-well plates. Cells were washed twice with PBS and resuspended in PBS for flow cytometry analysis. The geometric mean fluorescence intensity of each target population was determined after gating cells by forward and side scatter and selecting the full width at half maximum of the population in the corresponding fluorescence channel. A standard curve was generated using Quantum Simply Cellular anti-Mouse IgG beads stained with the same antibody used for the target cells. Molecules per cell were then calculated for each cell line from the standard curve and the geometric mean fluorescence intensity of the target-cell population. Antibodies used for antigen-density calibration included anti-HER2 APC, anti-CD19 APC, and anti-ROR1 APC, as listed in the flow cytometry antibody table.

**Primary human T cell isolation and culture**

Human primary T cells were isolated from de-identified donor Leukopaks obtained under institutional guidelines. Primary human CD4+, CD8+, and CD3+ T cells were isolated via negative selection (STEMCELL Technologies #17952, #17953, #17951). T cells were then cryopreserved at 10M-20M/mL in CELLBANKER 1 (Amsbio #11910). T cells were cultured in complete human T cell medium (HTCM) with X-VIVO 15 (Lonza #04-418Q), 5% human AB serum (Valley Biomedical #HP1022HI), 10 mM N-acetyl L-cysteine (Sigma #A9165-25G) neutralized with NaOH (Teknova #N4710), 55 uM beta-mercaptoethanol (Fisher 21985023), and supplemented with 30 units of IL-2 (BRB preclinical biologics repository).

**T cell viral transductions**

Lentivirus was produced by co-transfection of 70% confluent Lenti-X 293T cells (Takara #632180) with a transgene expression vector, pCMVdR8.91 and pMD2.G viral packaging plasmids, and polyethylenimine (PEI) transfection reagent (Kyfora Bio #24765-100). Primary T cells were thawed and plated at 1M/mL, and 24 hours later, activated with Dynabeads Human T-Activator CD3/CD28 for T Cell Expansion (ThermoFisher Scientific #11131D). After 24 hours of activation, T cells were virally transduced with 48-hour harvested lentivirus filtered through a 45 μm syringe filter for 24 hours. At Day 5 post T cell activation, Dynabeads were removed with pipet disruption and a magnet. T cells were stained and sorted on a BD FACSAria II (BD Biosciences) via positive myc-tag expression for surface transgene expression. For synNotch to CAR circuits, positive myc-synNotch expression and negative HA-CAR expression were selected for transgene-expressing and non-leaky T cells based on untransduced T cells. T cells were expanded for 11 to 15 days post sorting and cultured between 0.5M/mL – 1M/mL during expansion before *in vitro* or *in vivo* assays. Basal CAR expression reported in this study was measured after post-sort expansion and resting, as indicated in the figure legends.

***In vitro* T cell cytotoxicity assays with live cell imaging**

Primary human CD8+ or CD3+ T cells expressing a CAR or a synNotch-CAR circuit were co-cultured with cancer cell line targets expressing various levels of antigen (HER2, CD19, ROR1) for 72 hours in a 37°C, 5% CO_2_ incubator. In all assays, adherent cancer cells were seeded in their respective medium for adhesion 4-6 hours prior to the addition of T cells with complete human T cell media in 96-well flat bottom tissue culture plates at various effector to target cell ratios as indicated in the figure legends. Target cell lysis was measured by the target cell area and fluorescent marker intensity via live-cell imaging on the Incucyte SX5 (Sartorius). T cell proliferation and phenotype markers were assessed via flow cytometry.

**Mathematical calculations of killing**

On-target killing: % specific lysis of targets in co-cultures with high HER2 antigen-density cancer cells

$$On-target killing \%=\left( 1-\frac{High HER2 Target Cell Area with engineered T cells}{High HER2 Target Cell Area with untransduced T cells} \right)*100$$

Off-target killing: % specific lysis of targets in co-cultures with low HER2 antigen-density cancer cells

$$Off-target killing \%=\left( 1-\frac{Low HER2 Target Cell Area with engineered T cells}{Low HER2 Target Cell Area with un-engineered T cells} \right)*100$$

**Discrimination Score (DS):** Discrimination score was calculated as DS = (K_max_/100) × [1 − (K_min_/100)], where K_max_ and K_min_ are the percent target killing measured at high- and low-antigen density, respectively. DS= 0 being no discrimination and 1 being perfect discrimination of on-target and off-target cells.

$$DS =Ontarget killing *(1-OffTarget killing)$$

**Solution of the ODEs**

Modeling of the dynamic interactions between synNotch-CAR T cells and cancer target cells are described by a non-linear system of ordinary differential equations (ODEs). The state of the system is defined by three continuous time-dependent variables: cancer target cell population $C(t)$, total T-cell population $T(t)$, and the active CAR-expressing fraction of T cells $f_{\text{CAR}}\left( t \right)$. Given the parameters, basal fraction of CAR expressing cells, killing potency, proliferation limits, and initial cell seeding abundances ($C\left( 0 \right)=10,000$ cancer cells and $T\left( 0 \right)=30,000$ T cells for a 3:1 Effector-to-Target ratio), the non-linear system of ODEs is solved numerically in Python using the scipy.integrate.odeint routine with the LSODA numerical integration scheme. Relative and absolute error tolerances were set to $\text{rtol}={10}^{-6}$ and $\text{atol}={10}^{-8}$, respectively. Integrations were evaluated over a 72-hour time course ($t\in\left[ 0,72 \right]$ hours) discretized into 500 uniform time steps.

**Formulation of ODEs and Constitutive Kinetic Rates**

The rate of change of target cancer cells $C\left( t \right)$ is governed by baseline cancer proliferation and T-cell-mediated cytotoxicity:

$$\frac{dC}{dt}=r_{\text{tumor}}*C\left( t \right)-k_{\text{kill,eff}}\left( \rho,f_{\text{CAR}} \right)*C\left( t \right)*T\left( t \right)$$

where $r_{\text{tumor}}=0.00\text{ h}^{-1}$ during standard cytotoxicity assays. The effective killing rate $k_{\text{kill,eff}}$ is decoupled into baseline leak killing ($k_{\text{leak}}$) and antigen density-dependent induced CAR killing ($k_{\text{induced}}$):

$$k_{\text{leak}}=f_{\text{leak}}*k_{\text{kill,base}}$$

$$k_{\text{induced}}=\max\left( 0,f_{\text{CAR}}\left( t \right)-f_{\text{leak}} \right)*k_{\text{kill,base}}*\left[ \frac{\rho}{5\times{10}^{5}+\rho} \right]$$

$$k_{\text{kill,eff}}=k_{\text{leak}}+k_{\text{induced}}$$

The base killing rate constant is defined as:

$$k_{\text{kill,base}}=\left( 5.16\times{10}^{-6}\text{ cell}^{-1}\text{h}^{-1} \right)*P_{\text{potency}}$$

where $P_{\text{potency}}$ is the intrinsic single-cell cytotoxicity factor for each circuit construct.

The total T-cell population $T\left( t \right)$ expands according to antigen-driven logistic proliferation modulated by the instantaneous Effector-to-Target (E:T) ratio $r_{\text{ET}}=T\left( t \right)/C\left( 0 \right)$:

$$\frac{dT}{dt}=k_{\text{expansion}}*\text{activation}\left( \rho,f_{\text{CAR}},C \right)*T\left( t \right)*\left[ 1-\frac{T\left( t \right)}{T_{\text{max}}} \right]$$

$$T_{\text{max}}=M_{\text{exp}}*T\left( 0 \right)$$

$$\beta\left( E:T \right)=\left\{ \begin{matrix} 0.5+0.5*r_{\text{ET}} & \text{if }r_{\text{ET}}<1.0 \\ 1.0+\min\left( 1.0,\frac{r_{\text{ET}}-1.0}{2.0} \right) & \text{if }r_{\text{ET}}\geq1.0 \end{matrix} \right.$$

$$k_{\text{expansion}}=\alpha_{\text{base}}*\beta\left( E:T \right)$$

$$\text{activation}=f_{\text{CAR}}\left( t \right)*\left[ \frac{C\left( t \right)}{1000+C\left( t \right)} \right]*\min\left( 1.0,\frac{\rho}{{10}^{6}} \right)$$

The active CAR-expressing T-cell fraction $f_{\text{CAR}}\left( t \right)$ evolves between the baseline uninduced leak floor $f_{\text{leak}}$ and the fully induced CAR ceiling $f_{\text{max}}$:

$$\frac{df_{\text{CAR}}}{dt}=k_{\text{induction}}*\left[ f_{\text{target}}\left( \rho\right)-f_{\text{CAR}}\left( t \right) \right]*\left[ \frac{C\left( t \right)}{100+C\left( t \right)} \right]-k_{\text{decay}}*\left[ f_{\text{CAR}}\left( t \right)-f_{\text{leak}} \right]*\left[ 1-\frac{C\left( t \right)}{100+C\left( t \right)} \right]$$

$$a\left( \rho\right)=\text{clip}\left( \frac{\log_{10} \left( \rho\right)-4.0}{7.0-4.0},0,1 \right)$$

$$f_{\text{target}}\left( \rho\right)=f_{\text{leak}}+a\left( \rho\right)*\left( f_{\text{max}}-f_{\text{leak}} \right)$$

where $k_{\text{induction}}=0.10\text{ h}^{-1}$ and $k_{\text{decay}}=0.05\text{ h}^{-1}$.

**Parameter estimation**

We estimated model parameters $P_{\text{potency}}$(killing potency multiplier), $\alpha_{\text{base}}$ (baseline proliferation rate), and $M_{\text{exp}}$(maximum fold-expansion capacity cap) by fitting percentage target killing (% lysis) and Discrimination Score (DS) values obtained at day 3 (72 hours) post-incubation with target cells across E:T ratios (1:3, 1:1, and 3:1). Constitutive basal leak fraction $(f_{\text{leak}}$) and maximum CAR fractions ($f_{\text{max}}$) were fixed directly from flow cytometry measurements: $f_{\text{leak}}=6.0\%$ (HER2), $3.5$ (CD19), and $0.2$ (ROR1); $f_{\text{max}}=22.0\%$ (HER2), $22.0\%$ (CD19), and $20.0\%$ (ROR1).

We minimized the weighted multi-objective cost function below using the L-BFGS-B algorithm in Python (scipy.optimize.minimize routine with parameter bounds $P_{\text{potency}}\in\left[ 0.1,5.0 \right]$, $\alpha_{\text{base}}\in\left[ 0.0001,2.0 \right]$, and $M_{\text{exp}}\in\left[ 1.0,5.0 \right]):$

$$\text{Cost Function}=w_{\text{kill}}*\left[ 2*\left( \frac{K_{\text{min,expt}}-K_{\text{min,model}}}{100} \right)^{2}+\left( \frac{K_{\text{max,expt}}-K_{\text{max,model}}}{100} \right)^{2} \right]+\frac{1}{N_{\text{ET}}}*\sum_{E:T} \left( DS_{\text{expt}}-DS_{\text{model}} \right)^{2}+\mathcal{P}\left( T_{\text{max}} \right)$$

where $K_{\text{min}}$ and $K_{\text{max}}$ denote the percentage target killing at low (${10}^{4}$) and high (${10}^{7}$) antigen densities, respectively. Discrimination Score is defined as $DS=\left( K_{\text{max}}/100 \right)*\left( 1-K_{\text{min}}/100 \right)$. Off-Target killing values were set to $47.0\%$ (HER2), $29.0\%$ (CD19), and $0.0\%$ (ROR1); target maximal killing values were set to $96.0\%$ (HER2), $97.0\%$ (CD19), and $71.0\%$ (ROR1). The weighting factor was set to $w_{\text{kill}}=10.0$. $\mathcal{P}\left( T_{\text{max}} \right)$ is a penalty term enforcing an upper threshold of 90,000 total T cells by 72 h: $\mathcal{P}\left( T_{\text{max}} \right)=0$ if $T\left( 72\text{h} \right)\leq90,000$, and $\left[ \frac{T\left( 72\text{h} \right)-90,000}{10,000} \right]^{2}$ if $T\left( 72\text{h} \right)>90,000$. The fit parameters for the three circuits are in **Table 1**.

**Dose-response and sensitivity analysis**

Using the estimated parameters, dose-response killing curves were generated by logarithmically discretizing target antigen density into 30 points $\rho\in\left[ {10}^{4},{10}^{7} \right]$ molecules/cell and simulating target cell lysis at 72 hours. Circuit modifications (+mScarlet, +Degron, +mScarlet+Degron) were modeled by scaling baseline leak fractions according to empirical flow cytometry ratios: 3x leak reduction for +mScarlet, 4x leak reduction for +degron, and 12x leak reduction for dual +mSc+D.

Discrimination Score sensitivity across parameters was evaluated sweeping Leakiness (0%–15%), Killing Potency (0.1x–10x), Max CAR Fraction (5%–80%), and Proliferation Rate (0.0–1.0).

**SynNotch induction and CAR degradation kinetics via myc-bead activation assays**

To assess proliferation, primary human CD3+ T cells were stained with CellTrace dyes at 1:1000 for 30 mins at 37°C and washed with 10% FBS in PBS twice prior to bead activation assay. For synNotch induction and surface CAR expression measurements, 20,000 T cells were co-cultured with anti-myc magnetic beads (1:2000 dilution) (ThermoFisher #88843) in 96-well round bottom plates with a total volume of 200 μL of complete human T cell medium. The plate was centrifuged at 100 g for 1 min to promote bead and T cell interactions on the bottom of the plate. Beads were pipet disrupted and removed via magnetic plates at various timepoints as indicated in the figure legends. CAR expression, T cell proliferation, and T cell phenotype were assessed via flow cytometry (NovoCyte Quanteon).

**Flow cytometry and antibodies**

Novocyte Quanteon Flow Cytometer (Agilent) was used to assess T cell proliferation, receptor expression, and phenotype marker expression. All antibodies were used at 1:100 dilution in 10% FBS and PBS with 30 min staining at 4°C. At the end of staining, cells were washed with 10% FBS and PBS twice before flow cytometry analysis.

| **Target** | **Fluorochrome** | **Clone** | **Catalog** |
| --- | --- | --- | --- |
| myc | AF647 | 9B11 | CellSignaling 2233S |
| myc | PE | 9B11 | CellSignaling 3739S |
| HA | AF647 | 6E2 | CellSignaling 3444S |
| HA | AF488 | 6E2 | CellSignaling 2350S |
| CD4 | PerCP-eFluor710 | SK3 | Invitrogen 46-0047-41 |
| CD8 | FITC | SK1 | BioLegend 980908 |
| TCR α/β | BV421 | IP26 | BioLegend 306722 |
| CD45RA | BV785 | HI100 | BioLegend 304140 |
| CD62L | PE-Cy7 | DREG-56 | BioLegend 304822 |
| HER2 | APC | 24D2 | BioLegend 324408 |
| CD19 | APC | HIB19 | BioLegend 302212 |
| ROR1 | APC | 2A2 | BioLegend 357806 |
| Viability dye | Near-IR |  | Invitrogen L10119 |

**Immunofluorescence staining and fluorescence microscopy**

T cells were fixed in 4% paraformaldehyde in PBS for 10 min at room temperature. Following fixation, cells were washed three times with PBS and permeabilized with 0.5% Triton X-100 in PBS for 10 mins. Cells were then washed three times with PBS and blocked with 3% BSA in PBS for 1 hour at room temperature. Cells were then incubated with primary antibodies (Mouse anti-HA antibody, Thermo Fisher #26183, 1:400 dilution, Rabbit anti-Calnexin antibody, Thermo Fisher #PA5-34754, 1:1000 dilution) in blocking buffer overnight at 4°C. After washing three times with PBS, cells were incubated with secondary antibodies (Goat anti-Rabbit IgG (H+L) secondary antibody, AlexaFluor488, Thermo Fisher #A-11001, 1:500 dilution; Goat anti-Rabbit IgG (H+L) secondary antibody, AlexaFluor633, Thermo Fisher #A-21070, 1:500 dilution) and DAPI in blocking buffer for 30 min at room temperature. Cells were then washed three times with PBS and imaged. Fluorescence confocal microscopy was performed with a Zeiss AxioObserver microscope with 100X oil immersion objectives, outfitted with a Yokogawa spinning disk confocal head and Cascade II:512 camera.

***In vivo* mouse experiments**

All animal experiments were performed in accordance with protocols approved by the Stanford Administrative Panel on Laboratory Animal Care. Male and female NSG mice (JAX #005557) were bred by the Stanford VSC Breeding Colony Management Services (BCMS) core. For tumor xenograft experiments, 8- to 12- weeks-old animals were subcutaneously inoculated with engineered cancer cells in 50% PBS and 50% Matrigel (Corning #256255) at 1-3 M/flank as indicated per experiment. At 8-10 days post tumor inoculation, mice were randomized via RandoMice v1.1.7 by tumor size, sex, and weight, and 0.5-10M CD3+ T cells were injected intravenously in 200 μL of PBS with n = 5-8 mice per treatment group. Tumor volumes are monitored 2-3 times a week via caliper measurements until endpoints (loss of over 20% bodyweight, tumor volume exceeding 2000 mm^3^, immobility, and hunching).

**Tissue processing and Immunohistochemistry Staining.**

Mouse tumors were collected at the experimental endpoint and fixed in 4% paraformaldehyde in PBS overnight. Fixed tissues were washed in 70% ethanol and sent for FFPE embedding and tissue slide preparation by Histo-Tec Laboratory Inc. Following deparaffinization, heat-induced antigen retrieval was performed in Tris-EDTA pH=9.0. A peroxidase, avidin, biotin, and protein block was performed prior to primary antibody incubation (Agilent Dako #S2003, Vector Laboratories #SP-2001, Vector Laboratories #5030) for 20-30 mins sequentially. Slides were then stained with primary antibodies (Abcam ab16662, anti-ErbB2, 1:100 dilution, 200 uL) overnight at 4°C. The following day, slides were stained with secondary antibodies (Vector Laboratories #PK-4001, goat anti-rabbit, 1:500 dilution, 200 uL) for 45 mins. Slides were lastly stained with peroxidase for 30 mins (Vector Laboratories #SK-4100), DAB for 4 mins (Vector Laboratories #SK4400), and hematoxylin counterstaining. Completed slides were dehydrated and mounted for scanning and analysis using QuPath 0.5.1.

**Table 1: Calibrated Model Parameters:**

| **Circuit** | **Fixed Leak (%)** | **Potency (P)** | **Proliferation Base (α)** | **Max Expansion (M)** | **Simulated K_min_**  **(off-Target)** | **Simulated K_max_**  **(on-Target)** | **Discrimination**  **Score (DS)** |
| --- | --- | --- | --- | --- | --- | --- | --- |
| HER2-HER2 | 6.00% | 0.94 | 1.0000 | 3.00× | 47.5% (47.0%) | 99.5% (96.0%) | 0.52 |
| HER2-CD19 | 3.50% | 0.89 | 1.0000 | 3.00× | 29.7% (29.0%) | 99.3% (97.0%) | 0.70 |
| HER2-ROR1 | 0.20% | 0.69 | 0.0010 | 1.80× | 1.5% (0.0%) | 71.6% (71.0%) | 0.71 |
